# Cell state-specific metabolic networks govern ferroptosis versus apoptosis in small cell lung cancer

**DOI:** 10.64898/2026.03.27.714827

**Authors:** Jun W. Kim, Christina M. Bebber, Yuqin Dai, Sarah Bopp, Anna Edenhofer, Albert M. Li, Thomas Rösner, Lena Berning, Ming Yang, Logan B. Leak, Jenny Stroh, Bindesh Shrestha, Ali T. Abdallah, Dimitrios Prymidis, Hernando J. Olivos, Maya Baron, Thuyen Nguyen, Yan Ting Shue, Yoko Nishiga, Alexandros P. Drainas, Andrea C. Chaikovsky, Krystina Szylo, Yang Li, Yun P. Kang, Parvathy Manoj, Alvaro Quintanal Villalonga, Charles M. Rudin, Gina M. DeNicola, Scott J. Dixon, Christian Frezza, Jiangbin Ye, Silvia von Karstedt, Julien Sage

## Abstract

Cellular heterogeneity and plasticity are hallmarks of cancer that contribute to tumor growth and therapy resistance. Here we investigated metabolic heterogeneity in small cell lung cancer (SCLC), an aggressive neuroendocrine (NE) cancer type. Through integrated transcriptomic and metabolomic analyses, we identified a universal dependency on exogenous cysteine/cystine (Cys) across all NE/non-NE SCLC cell states. Notably, NE and non-NE cells with low levels of the ASCL1 transcription factor die from ferroptosis upon Cys depletion. In contrast, ASCL1-high cells die from apoptosis but are ferroptosis resistant. This resistance to ferroptosis is driven by the direct upregulation of the gene coding for the GCH1 enzyme by ASCL1, which results in higher levels of the BH4/BH2 antioxidants. Accordingly, combining cysteine depletion with BH4/BH2 synthesis inhibition effectively reduces tumor growth in patient-derived xenografts. This work elucidates distinct metabolic states in SCLC and suggests new approaches to induce cell death in this lethal form of cancer.

## INTRODUCTION

Cellular heterogeneity and plasticity are critical traits of tumor evolution and resistance to therapy^1,2^. Small cell lung cancer (SCLC), a paradigm for the role of tumoral heterogeneity in resistance to treatment^3-9^, undergoes rampant plasticity between cellular states defined by transcription factors regulating neuroendocrine (NE) differentiation. Two major NE subtypes exist in SCLC, SCLC-A and SCLC-N, where cancer cells exhibit generally mutually exclusive high expression of ASCL1 and NEUROD1, respectively^9-12^. However, intratumoral heterogeneity can be prominent, with individual SCLC tumors harboring different types of NE cells as well as less/non-neuroendocrine (non-NE) cancer cells^3,6,7,12-15^. Non-NE (ASCL1^low^ and NEUROD1^low^) cancer cells in tumors are thought to be genetically similar to NE cancer cells but can emerge from ASCL1^high^ or NEUROD1^high^ NE cancer cell populations by epigenetic reprogramming, including upon activation of Notch and YAP1 signalling^3,16-20^. While transcriptional subtype-specific targeting approaches have been proposed for SCLC^5,8,9,21-25^, accumulating evidence of plasticity between cell states in SCLC^7,15,26-31^ underscores the importance of targeting SCLC cells across all cell states for effective treatment.

Cancer cells have altered metabolism^32,33^, and the distinct metabolic features of cancer cells create potential metabolic vulnerabilities that can be exploited for therapeutic intervention^34,35^. Accumulating evidence also points to inter- and intratumoral metabolic heterogeneity, the origins of which are still poorly understood^36-41^. A major goal of the field is to characterize cell type- and cell state-dependent metabolic heterogeneity to both gain both a deeper understanding of cancer metabolism and use this knowledge for more effective therapeutic strategies against cancer. Metabolic vulnerabilities have begun to emerge for in SCLC^22,42^, but an unresolved issue in the field is how changes in cell state influence cellular metabolism and whether such metabolic shifts could expose targetable bottlenecks in SCLC.

In the case of fast-growing tumors such as SCLC, induction of cancer cell death is critical to inhibit tumor growth. While DNA damage is known to trigger p53-dependent intrinsic apoptosis via mitochondrial activation of the caspase cascade^43^, the majority of treatment naïve SCLC already present with bi-allelic inactivation of p53 (and RB) ^44^. Thus, the p53-dependent intrinsic apoptosis pathway is likely already disabled in SCLC tumors before standard-of-care chemotherapy or radiation therapy. Notably, DNA-damage causing therapeutic agents can also trigger caspase-dependent cell death via the accumulation of reactive oxygen species (ROS)^45^, which can bypass the requirement for p53. Ferroptosis, a caspase-independent cell death critically dependent on ferrous iron^46^, is characterized by excessive lipid peroxidation driven by lipid ROS and ensuing loss of plasma membrane integrity and osmotic cytoplasmic swelling^47-49^. Glutathione peroxidase 4 (GPX4) constitutively hydrolyses lipid hydroperoxides and thereby protects cells from ferroptosis^50^. GPX4 requires glutathione (GSH) as an electron donor to reduce lipid hydroperoxides. Consequently, impairment of the two major routes of cysteine supply for GSH synthesis, cystine import from the extracellular space via the cystine/glutamate antiporter System xc, and *de novo* synthesis via the transsulfuration pathway, can trigger ferroptosis^51^. Notably, a lipid composition rich in ether-linked polyunsaturated fatty acids (PUFAs) was shown to cause ferroptosis sensitivity of non-NE SCLC^21^. The lipid synthesis pathway was shown to be required for optimal SCLC growth^52^, suggesting additional metabolic bottlenecks.

Here, using integrated transcriptomic and metabolomic approaches, we found that SCLC cells in all states are dependent on exogenous cysteine/cystine (Cys) for their survival. We also uncovered that SCLC cells in the ASCL1^high^ state are resistant to the ferroptosis that is normally induced by Cys depletion and we identified an ASCL1-GCH1-BH4/2 axis that protects these cells from ferroptosis. These observations suggest various combination therapies, which, together with Cys depletion, can potently inhibit the growth of SCLC tumors from distinct subtypes.

## RESULTS

### Integrated metabolomic and transcriptomic analyses in mouse models of SCLC reveal a decrease in the glutathione pathway in the ASCL1^high^ neuroendocrine state

To investigate possible metabolic changes associated with distinct SCLC cell states, we performed an unbiased analysis of differentially regulated metabolites and transcripts in two complementary genetically engineered mouse models of SCLC. First, we used the *Rb^flox/flox^*;*p53^flox/flox^*;*Rbl2^flox/^*^flox^;*Hes1^GFP^*(*RPR2;Hes1^GFP^*) model in which GFP expression can be used to isolate NE (GFP^neg^, ASCL1^high^) and non-NE SCLC cells (GFP^high^, ASCL1^low^, with higher Notch and YAP1 signaling) from tumors^16^ (**Figure 1A**). Second, we isolated cancer cell populations from the *Rb^flox/flox^*;*p53^flox/flox^ (RP)* mouse model of SCLC^53^ based on their growth in suspension (NE ASCL1^high^ cells, ‘floaters’) or their attachment to cell culture plates (non-NE, ASCL1^low^ and YAP1^high^, ‘stickers’)^3,21^ (**Figure 1A**).

**Figure 1:**
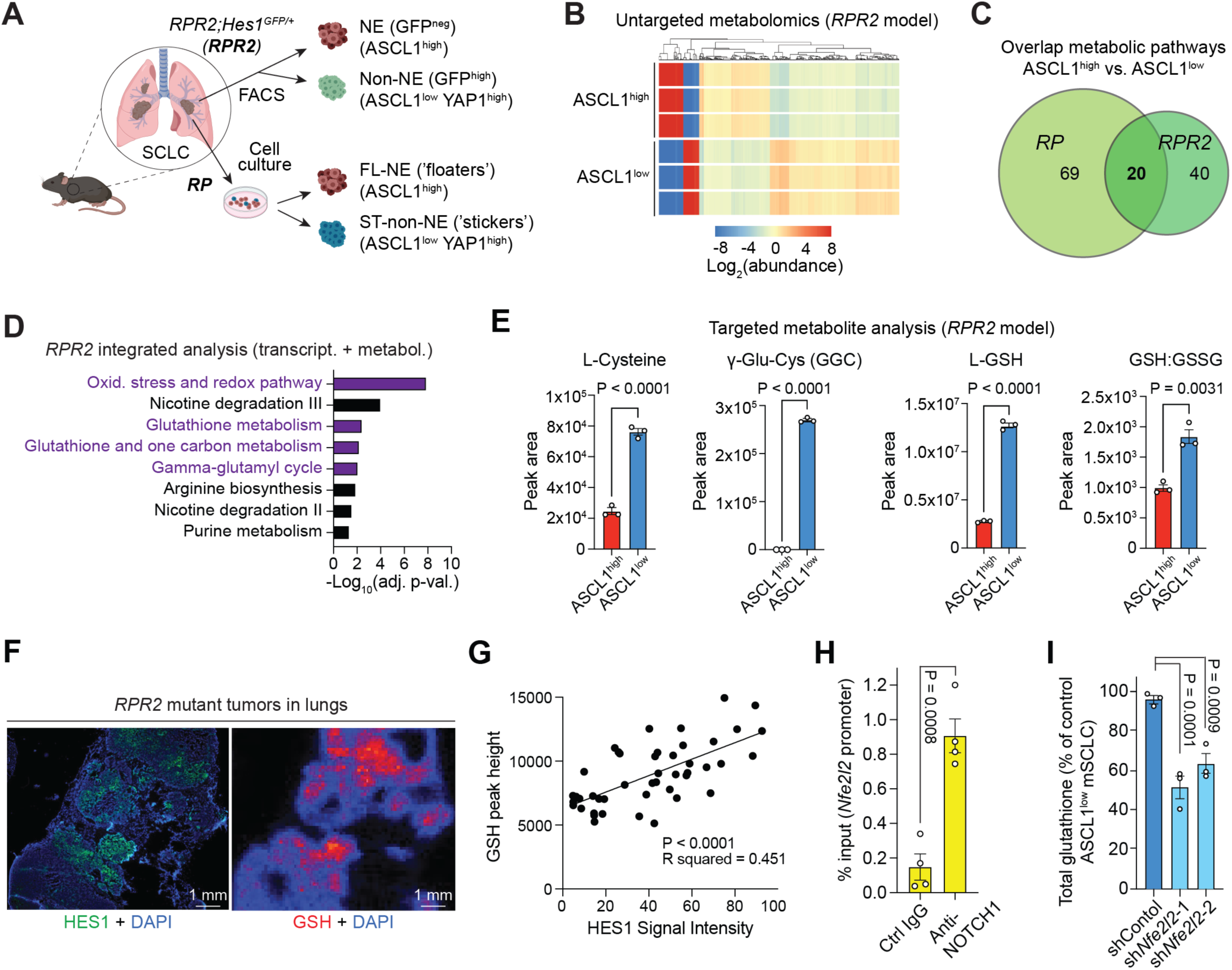
Integrated metabolomic and transcriptomic analysis reveals differential regulation of glutathione in distinct SCLC neuroendocrine cell states. **A.** Schematic of the two SCLC mouse models: the *RPR2;Hes1^GFP^*model (*RPR2*) (*Rb1^Δ/Δ^;p53^Δ/Δ^;Rbl2^Δ/Δ^;Hes1G^FP/+^*mutant tumors), in which non-neuroendocrine (non-NE, ASCL1^low^ YAP1^high^) SCLC cells can be separated from NE ASCL1^high^ SCLC cells using GFP expression, and the *RP* model (*Rb1^Δ/Δ^;p53^Δ/Δ^*), in which non-NE “stickers” (ST, ASCL1^low^ YAP1^high^) can be separated from NE “floaters” (FL, ASCL1^high^) in culture. **B.** Hierarchical clustering analysis of statistically significant (P<0.005) metabolites with a fold-change of at least 2 in the *RPR2* model comparing ASCL1^high^ and ASCL1^low^ SCLC cells. **C.** Venn diagram showing overlapping pathways identified in overrepresentation analysis of untargeted metabolomics in the two mouse models as in (A). See Tables S1,2. **D.** Selected pathways differentially regulated transcriptionally (adjusted P-value>0.05, n=4) and metabolically (with n≥3 matched metabolites) in the *RPR2* model comparing ASCL1^high^ and ASCL1^low^ SCLC cells. Purple: glutathione-associated pathways. **E.** Relative abundance (peak area) of selected metabolites detected by LC/MS in ASCL1^high^ and ASCL1^low^ SCLC cells (n=3) from the *RPR2* model. **F.** Representative fluorescent imaging (left) of HES1 (anti-HES1, green) and mass spectrometry imaging (right) of GSH (red) in a representative section from *RPR2* mutant mouse lungs. DAPI marks the DNA in blue. Scale bar, 1 mm. **G.** Average peak area (5 × 5 pixels) of GSH and HES1 fluorescence signal intensity in randomly chosen tumor regions as in (F) (n=45 points, n=2 *RPR2* mutant mouse lungs). **H.** ASCL1^high^ mouse SCLC cells were transduced with a lentiviral vector expressing the NOTCH1 intracellular domain (N1ICD) and chromatin immunoprecipitation (ChIP) was performed, followed by qPCR around a consensus sequence for RBP-J binding (the partner of NICD) in the *Nfe2l2* promoter (n=4). **I.** Average levels of total glutathione (GSH) in ASCL1^high^ and ASCL1^low^ mouse SCLC cells (mSCLC) quantified as percent of glutathione colorimetric assay readout in control wild-type cells. Two independent shRNAs were used to knock-down *Nfe2l2* (coding for NRF2) (n=3). The P-values for (D) were calculated from pathway analysis using the RNA-seq data. P-values for (E) and (H) were calculated by the unpaired Student’s t-test. The P-value for (G) was calculated using the F-test. Sidak test following one-way analysis of variance (ANOVA) were performed in (I) (P<0.0001). Error bars indicate mean ± SEM.

Bulk metabolomic analyses were performed comparing ASCL1^high^ and ASCL1^low^ cancer cells in both models. In cells derived from *RPR2* mutant tumors, 1296 metabolites were detected, 474 of which were significantly different by greater than 2-fold between the ASCL1^high^ and ASCL1^low^ populations (**Figure 1B** and **Figure S1A**). Similarly, metabolomics of the *RP* mutant SCLC model detected 224 metabolites, 187 of which were significantly different between ASCL1^high^ ‘floaters’ and ASCL1^low^ ‘stickers’ (**Figure S1B,C**). Accordingly, ASCL1^high^ and ASCL1^low^ SCLC cells in both models clustered separately by principal component analysis (PCA) (**Figure S1D,E**). Pathway analysis in the *RPR2* model identified metabolic pathways impacted by at least three or more statistically significant metabolites between the two cell states, including in pathways related to purine metabolism, tRNA charging, oxidative stress and redox, and sphingolipid metabolism (**Table S1**). Notably, 20 of these overlapped with pathways significantly different between ASCL1^high^ ‘floaters’ and ASCL1^low^ ‘stickers’ in the *RP* model (**Figure 1C** and **Table S2**), highlighting the similarities between the two models.

Integration of these metabolomic data with RNA sequencing (RNA-seq) data^16^ further identified metabolic pathways associated with each cell state in the *RPR2* model (**Table S3**), with some of the top-ranked pathways related to glutathione-mediated redox balance (**Figure 1D**). A metabolic network generated using the genes and metabolites in the pathways around cysteine and glutathione in the *RPR2* model showed that ASCL1^high^ SCLC cells had decreased gene expression and metabolites involved in glutathione synthesis and utilization compared to ASCL1^low^ SCLC cells (**Figure S1F**). As the cysteine-glutathione metabolic pathway is critical for many cells, including cancer cells, to respond to oxidative stress^54-56^, we further investigated this observation.

The differences in key metabolites in the glutathione pathway (L-cysteine, γ-glutamylcysteine [GGC], and reduced L-glutathione [GSH]) between ASCL1^high^ and ASCL1^low^ SCLC cells were confirmed in both models by LC/MS analysis using metabolite standards (**Figure 1D** and **Figure S1G**). Moreover, the GSH to oxidized L-glutathione (GSSG) ratio, a measure for reducing capacity, was decreased in *RPR2-*derived ASCL1^high^ SCLC cells compared to ASCL1^low^ SCLC cells from the same tumors (**Figure 1E** and **Figure S1H,I**). Levels of the mRNA coding for cysteine dioxygenase type 1 (*CDO1*) and of downstream metabolites (hypotaurine, taurine, and pyruvic acid) of the sulfinic acid pathway, which is known to compete with the glutathione synthesis pathway for cysteine, were upregulated in ASCL1^high^ compared to ASCL1^low^ SCLC cells (**Figure S1F,J,K**). *In situ* metabolomics on sections from *RPR2* mutant tumors using desorption electrospray ionization (DESI) confirmed local enrichment of HES1^high^ (ASCL1^low^) SCLC cells and increased levels of glutathione within intact SCLC tumors (**Figure 1F,G** and **Figure S2A-C**).

Our experiments above raised the question how the ASCL1^low^ state sustains higher levels of GSH compared to the ASCL1^high^ state. A major difference between these two states is activation of the Notch signaling pathway in ASCL1^low^ cells^3,16,18,19^. Notch pathway activity was shown in another context to induce the activity of the NRF2 (Nuclear factor erythroid 2-related factor 2) antioxidant pathway^57^. Through chromatin immunoprecipitation (ChIP), we found that the transcriptionally active form of NOTCH1 (NOTCH1 intracellular domain, N1ICD) binds to regulatory regions of the *Nfe2l2* gene, which codes for NRF2, when N1ICD was ectopically expressed in ASCL1^high^ SCLC cells^3^ (**Figure 1H**). Accordingly, the expression of *Nfe2l2* and the known NRF2 target Glutamate–cysteine ligase catalytic subunit (*Gclc*), coding for the rate-limiting enzyme in the synthesis of GSH, was higher in *RPR2*- and *RP*-derived ASCL1^low^ compared to ASCL1^high^ SCLC cells in the RNA-seq analysis (**Figure S2D,E**). The analysis of single cell RNA-seq (scRNA-seq) data from SCLC cells undergoing an NE-to-non-NE transition *ex vivo* in the *RPM* mouse model (*Rb/p53/Myc*^15^) also revealed a negative correlation between *Ascl1* and NRF2 target genes, and a positive correlation with *Yap*1 (**Figure S2F**). Similar observations were made in human tumors (**Figure S2G**). NRF2 knock-down in ASCL1^low^ mouse SCLC cells in culture was sufficient to decrease the levels of GSH (**Figure 1H** and **Figure S2H**), indicating that elevated GSH levels are sustained at least in part through NRF2-mediated signaling in this cell state.

Altogether, using two genetically engineered mouse models of SCLC, we identified cell-state-dependent metabolic differences in redox state within heterogeneous tumors.

### SCLC cells exhibit cysteine dependence across cell states

Given that ASCL1^high^ SCLC cells showed decreased levels of cysteine and GSH, we hypothesized that mouse ASCL1^high^ SCLC cell lines may be particularly dependent on cysteine. To test this hypothesis, we selectively depleted Cys, as well as other amino acids, in the culture medium. Among the non-essential amino acids, Cys and glutamine were most critically required for cell viability (**Figure 2A** and **Figure S3A**). ASCL1^high^ cells showed a stronger dependency on exogenous Cys compared to ASCL1^low^ SCLC cells (**Figure 2B**). Importantly, NRF2 knock-down in ASCL1^low^ cells was sufficient to sensitize these cells to Cys depletion comparable to ASCL1^high^ cells (**Figure 2C** and **Figure S3B**). GSH supplementation also rescued cell viability loss upon Cys withdrawal (**Figure 2D** and **Figure S3C**). Moreover, Cys depletion was accompanied with accumulation of cellular reactive oxygen species (ROS) (**Figure 2E**).

**Figure 2:**
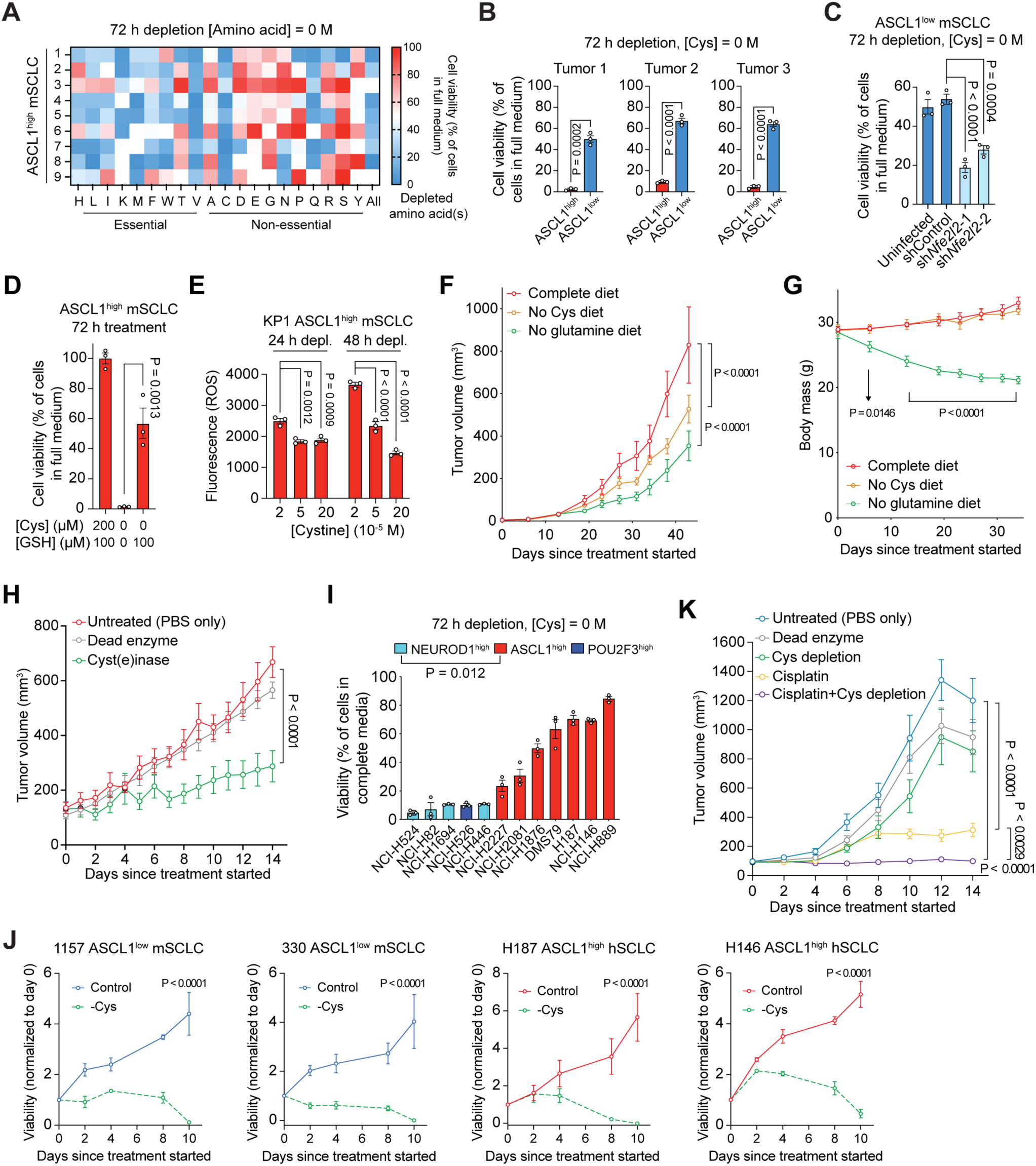
SCLC cells are dependent on exogenous Cys. **A.** Average viability (AlamarBlue assay) of ASCL1^high^ mouse SCLC (mSCLC) cell lines generated from independent *RPR2* mutant tumors and cultured in amino acid-depleted media for 72 h (n=3). **B.** Average viability (AlamarBlue assay) for pairs of ASCL1^high^ and ASCL1^low^ mouse SCLC cells isolated from *RPR2* mutant tumors 72 h after complete depletion of extracellular Cys (n=3). **C.** Average viability (AlamarBlue assay) of ASCL1^low^ mSCLC cells with stable knock-down of *Nfe2l2* (sh*Nfe2l2*) (coding for NRF2) or GFP (as a control, shControl) and cultured in Cys-depleted medium for 72 h (n=3). **D.** Average viability (AlamarBlue assay) of ASCL1^high^ mSCLC cells supplemented with GSH [100 µM] and cultured in complete medium [Cys = 200 µM] or Cys-depleted medium for 72 h (n=3).**D.** Measurement of reactive oxygen species (ROS) (with 2’,7’-dichlorodihydrofluorescein diacetate, DCFH-DA) in ASCL1^high^ mSCLC cells (KP1 cell line) cultured for 24 h or 48 h under depletion of extracellular Cys (n=3). Cells cultured in complete medium [Cys = 200 µM] were used as controls. Values are raw fluorescence values with background subtracted. **F.** Volumes of ASCL1^high^ SCLC allografts in mice on complete amino acid diet, no Cys diet, and no glutamine diet (n=10, 43 days). **G.** Body mass for mice with ASCL1^high^ SCLC allografts as in (F). **H.** Volumes of ASCL1^high^ SCLC allografts in mice treated with cyst(e)inase or dead enzyme (50 mg/kg, q.2.d., i.p.)for 14 days (n=6). **I.** Average viability (measured by AlamarBlue assay) of human cell lines from different SCLC subtypes cultured in Cys-depleted medium for 72 h. Numbers were quantified as percent of assay readout in cells in complete RPMI medium (n=3). **J.** Growth curves (AlamarBlue assay) of two ASCL1^low^ mSCLC cell lines and two human SCLC (hSCLC) ASCL1^high^ cell lines growing in full medium or Cys-depleted medium for 8 days (n=3). **K.** Growth of NCI-H82 human SCLC xenografts (NEUROD1^high^) treated with dead enzyme (50 mg/kg, q.2.d., i.p.) and Cys-depleted diet, cyst(e)inase (50 mg/kg, q.2.d., i.p.) and Cys-depleted diet (No Cys), cisplatin (4 mg/kg, b.i.w., i.p.), or No Cys and cisplatin (n=14). For (B), a Student’s t-test was used to calculate p-values. Error bars indicate mean ± SD. Sidak tests following one-way analysis of variance (ANOVA) were performed in (C) (P<0.0001), (D) (P<0.0001), (E) (P=0.0008), and (F) (P<0.0001). P-values were calculated by two-way analysis of variance (ANOVA) (P<0.0001) followed by the unpaired Student’s t-tests in (H) and (K) (P<0.0001). Error bars for all other figures indicate mean ± SEM.

Therapeutic Cys depletion can inhibit tumor growth in other cancer models^58-62^ and can enhance the effects of treatments that increase oxidative stress^45,63^. Indeed, we found moderate anti-tumor activity in mice bearing subcutaneous ASCL1^high^ mouse SCLC tumors fed with a Cys-depleted chow (**Figure 2F**), without any body mass loss (**Figure 2F**). In contrast, tumor inhibition in mice fed glutamine-deficient chow was accompanied by body mass loss (confirming a key role for glutamine in SCLC^64^) (**Figure 2F,G**).

Apart from dietary approaches, treatment with a cysteine-degrading enzyme can effectively break down and deplete extracellular Cys^65^. While wild-type human and mouse cystathionine gamma lyases (CGLs, cyst(e)inases) both potently inhibited the viability of mouse ASCL1^high^ SCLC cells, we found that engineered human CGL was significantly more potent in culture (**Figure S4A-E**). Strikingly, treatment of mouse ASCL1^high^ SCLC allografts with human pegylated CGL (PEG-cyst(e)inase) led to a significant anti-tumor response, as opposed to the dead enzyme control, without body mass loss (**Figure 2H** and **Figure S4F,G**).

Importantly, we could confirm Cys dependence also for a panel of human SCLC cell lines (**Figure 2I**). We noted, however, that acute Cys deprivation in NEUROD1^high^ cell lines nearly eliminated these cells, while, as a group, ASCL1^high^ cell lines showed higher residual viability in the short time frame of the assay (**Figure 2I**). Human ASCL1^high^ cell lines, as well as murine ASCL1^low^ cells, eventually also succumbed to cysteine depletion in longer depletion assays (**Figure 2J**). The one human POU2F3^high^ cell line tested also showed high dependency on exogenous Cys, similar to NEUROD1^high^ cells (**Figure 2I**), suggesting a varying but strong dependence on Cys across various cellular states in human SCLC.

Cisplatin is a standard-of-care chemotherapy agent in SCLC, and cisplatin treatment has been shown to induce oxidative stress associated with decreased levels of GSH^66^. We reasoned that in SCLC cell models highly sensitive to Cys depletion (i.e., NEUROD1^high^ cells), the anti-tumor effects of cisplatin treatment may be enhanced by Cys depletion. The cisplatin dosing regimen we used effectively decreased tumor volume for NCI-H82 xenografts, with some toxicity, as would be expected; combining cisplatin with Cys depletion using the PEG-cyst(e)inase further inhibited tumor growth, without additional loss of body mass (**Figure 2K** and **Figure S4H-J**).

Thus, cysteine dependence is a metabolic vulnerability in SCLC, and Cys depletion can enhance the therapeutic benefits of chemotherapy in this tumor type.

### Impaired transmethylation in all SCLC cell states creates a dependence on cysteine

Cells can obtain cysteine either via uptake of Cys from the extracellular space followed by its reduction to cysteine or through *de novo* synthesis via the transmethylation pathway, which normally converts methionine into homocysteine that can be used to generate glutathione (**Figure 3A**). The fact that SCLC cells across various cellular states were dependent upon Cys supplementation suggested restrictions to cysteine *de novo* synthesis. Indeed, when we examined the expression of key enzymes in these pathways in human SCLC cell lines^67^, we found that expression of the transsulfuration pathway enzyme Glycine N-methyltransferase (GNMT), which converts homocysteine into cysteine, was almost undetectable (**Figure 3B**). Low GNMT expression has been linked with Cys dependency in a few other cancer cell lines^68^. Using liquid chromatography-mass spectrometry (LC-MS) with [3-^13^C] serine (^13^C on 3^rd^ position in serine) as a tracer for transsulfuration pathway activity^69^ (**Figure 3C,D** and **Table S4**), we could not detect any activity in the SCLC cell lines tested, with no tracing detected towards L-cysteine or GSH (**Figure 3E,F**). Consistent with prior studies in a different context^67^, homocysteine rescued the growth of the majority of the Cys-dependent SCLC cell lines, while the upstream precursors could not (**Figure 3G**). Treating Cys-deprived NCI-H82 cells in medium containing [3-^13^C] serine with homocysteine led to rapid internalization of homocysteine and [3-^13^C] serine and ^13^C-labeling of downstream pathway members and a substantial decrease of total serine pool, reflecting serine consumption through the transsulfuration pathway (**Figure S5A-E**). Importantly, incubating Cys-deprived NCI-H82 cells expressing ectopic GNMT (**Figure 3B**) with [3-^13^C] serine resulted in increased [3-^13^C] labelling of GSH and GSSG and a decreased serine pool (**Figure 3H** and **Figure S5F,G**), indicative of reactivation of the transmethylation pathway. Consequently, ectopic GNMT expression was sufficient to fully support the proliferation of Cys-deprived NCI-H82 cells (**Figure 2I**). In a proteogenomic SCLC dataset with adjacent normal lung tissue^70^, SCLC tumors expressed comparable low levels of *GNMT* relative to adjacent normal tissue, but had higher expression of the cystine glutamate antiporter xCT (*SLC7A11*) (**Figure 2J**), which serves as a route to obtain cystine from the microenvironment and can support the viability of SCLC cells^21^.

**Figure 3:**
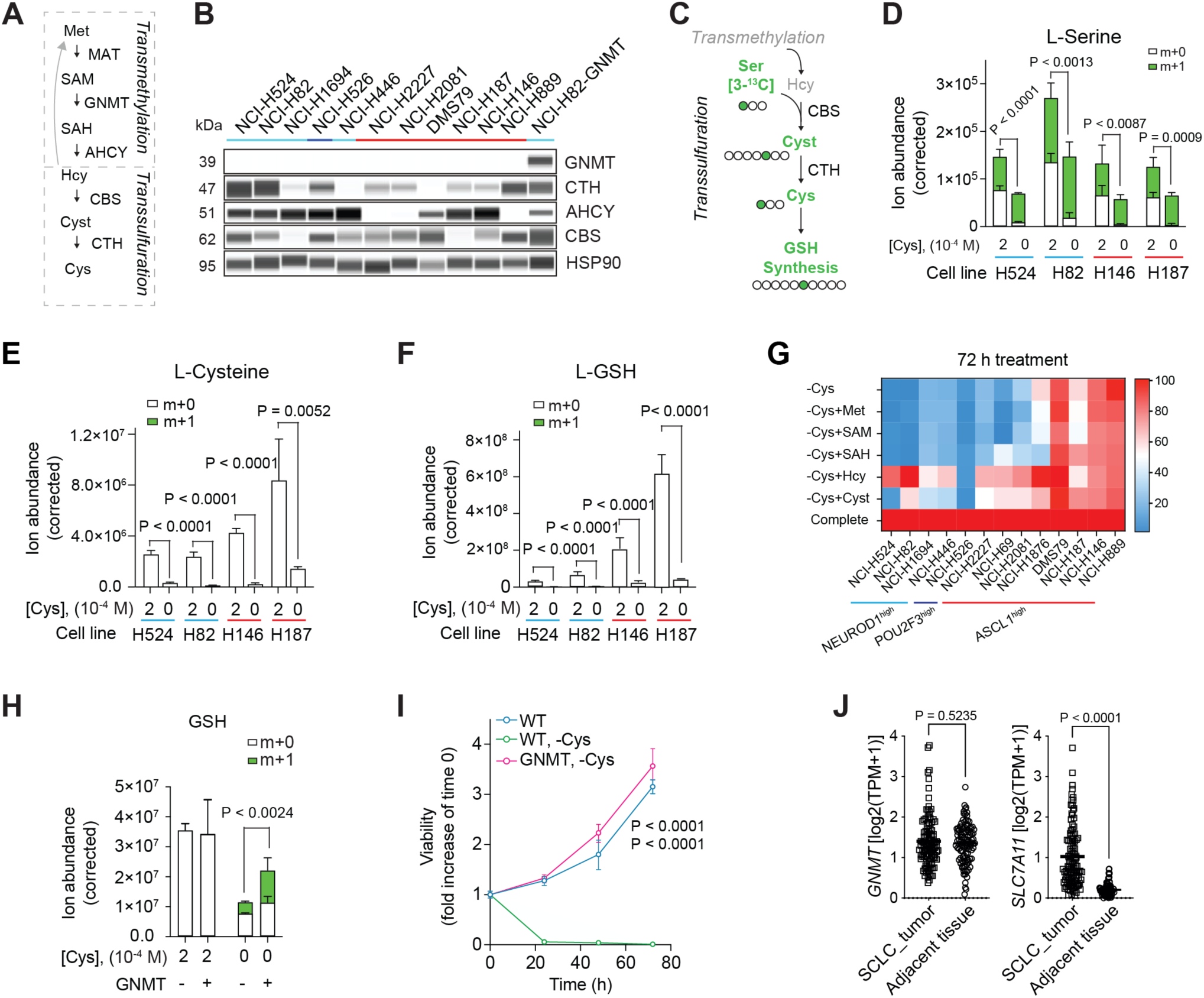
SCLC cells are dependent on exogenous Cys. **A.** Schematic for the *de novo* cysteine synthesis pathway. Met, methionine; SAM, S-adenosylmethionine; SAH, S-adenosylhomocysteine; Hcy, homocysteine; Cyst, cystathionine. **B.** Representative immunoassay for four enzymes as in (K) in human SCLC cell lines as in (I) (n=3). NCI-H82-GNMT cells are stably infected with a lentiviral vector expressing GNMT. HSP90 was used as a loading control. **C.** Schematic of [3-^13^C] serine isotope tracing. Clear circles indicate unlabeled carbon atoms. **D-F.** Labeled and unlabeled L-serine (D), L-cysteine (E), and reduced glutathione (GSH) (F) in NEUROD1^high^ and ASCL1^high^ human SCLC cell lines as in (B) in Cys-replete or Cys-deprived medium containing ^13^C-labeled L-serine (12 h, n=3). **G.** Average viability (AlamarBlue assay) in human SCLC cell lines as in (B) cultured in Cys-depleted medium supplemented with the indicated metabolic precursors of Cys [100 µM] for 72 h (n=3). **H.** Labeled and unlabeled GSH in NCI-H82 SCLC cells (NEUROD1^high^) ectopically expressing GNMT in Cys replete and depleted medium (n=4). **I.** Growth curves (AlamarBlue assay) of control NCI-H82 cells (WT) and NCI-H82 cells ectopically expressing GNMT as in (B) in full or Cys-depleted medium (n=3). **J.** *GNMT* and *SLC7A11* expression levels from an RNA-seq dataset (as in ^70^), comparing tumors and adjacent lung tissue (n=107) plotted as log2 of transcript per million (TPM)+1. P-values were calculated by the unpaired Student’s t-test. Error bars indicate mean ± SEM.

Collectively, these data indicate that SCLC cells are highly dependent on cysteine across various cellular states due to dysfunctional transmethylation for the synthesis of the central antioxidant GSH.

### Cys depletion triggers ferroptosis in ASCL1^low^ SCLC cells

Apart from serving as a direct radical-trapping agent, GSH can serve as a co-factor for glutathione peroxidase 4 (GPX4) (and other GPX enzymes). GPX4 is a central enzyme that protects against ferroptosis^47^, an iron-dependent type of cell death driven by excessive membrane lipid peroxidation^50^. GPX4 inhibition can selectively kill ASCL1^low^ SCLC cells^21^. Therefore, we next tested whether Cys depletion would induce ferroptosis in SCLC cells.

While cell death induced by Cys depletion was indeed rescued by the lipophilic radial trapping antioxidant ferrostatin-1 (Fer-1) in NEUROD1^high^ human and *RP*-mouse-derived ASCL1^low^ cell lines, this was not the case for human ASCL1^high^ and *RP*-mouse-derived ASCL1^high^ cell lines (**Figure 4A,B**). In support of the induction of ferroptosis in human NEUROD1^high^ and POU2F3^high^ cells, and in mouse ASCL1^low^ cells, lipid peroxides were significantly increased upon Cys depletion in these cellular states (**Figure 4C,D**) and reverted by co-treatment with Fer-1, but not by co-treatment with other cell death inhibitors, such as caspase inhibitors (Emricasan and QVD) and a RIPK1 inhibitor (Nec1s), blocking apoptosis and necroptosis, respectively (**Figure 4E** and **Figure S6A-C**). Of note, NEUROD1^high^ SCLC cells derived from the *RPM* mouse model^8^ were also highly sensitive to Cys depletion-induced ferroptosis (**Figure S6D,E**).

**Figure 4:**
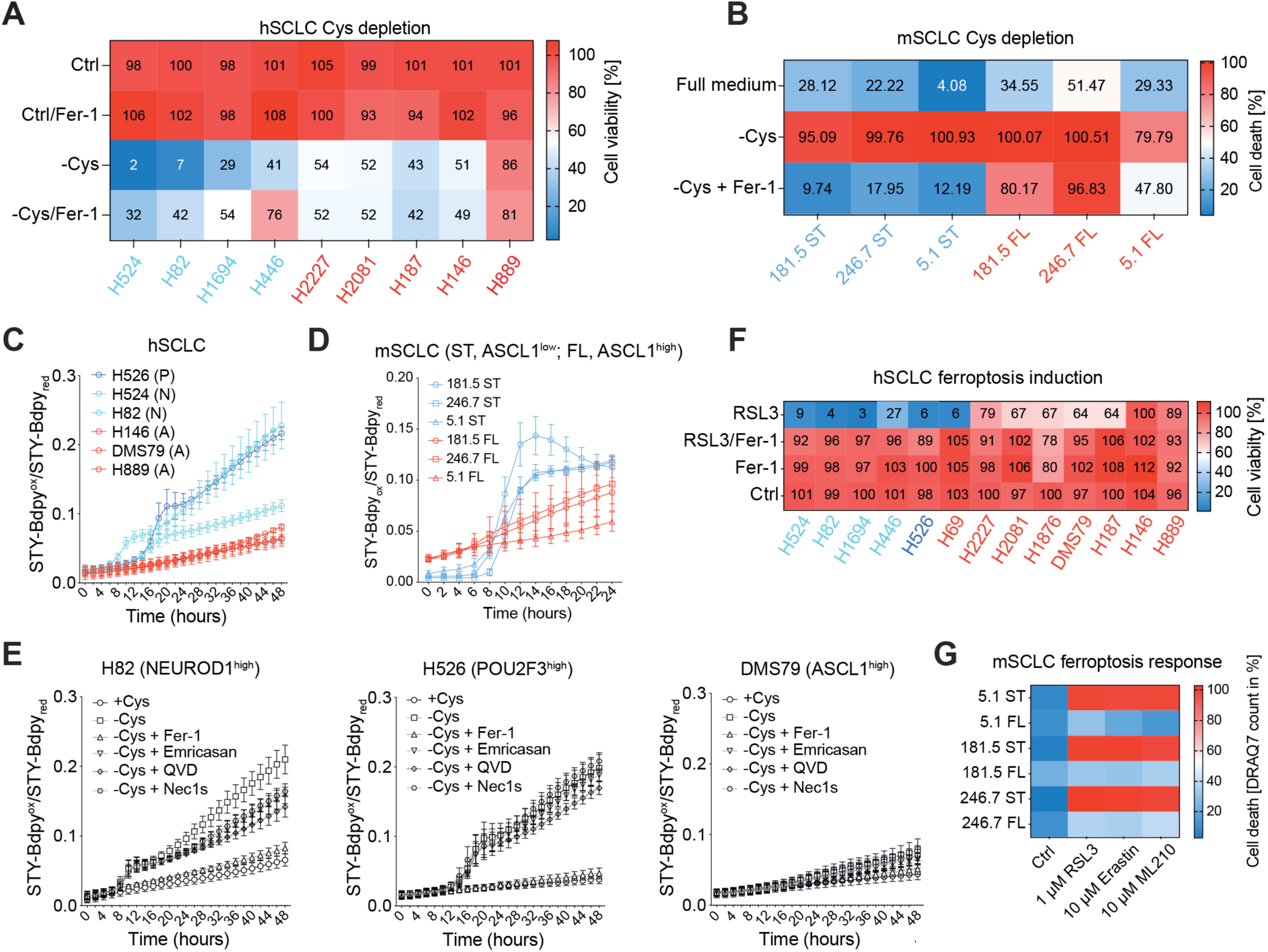
ASCL1^low^ SCLC cells undergo ferroptosis in response to Cys deprivation. **A.** Average viability (AlamarBlue assay) of four human NEUROD1^high^ and five human ASCL1^high^ SCLC cell lines (hSCLC) cultured in full medium, supplemented with ferrostatin-1 (Fer-1, a ferroptosis inhibitor) [1 µM], Cys depleted medium, or Cys depleted medium with Fer-1 for 72 h (n=3). **B.** Average cell death (DRAQ7 count/confluence in %) of *RP*-derived mouse ASCL1^high^ (FL) and ASCL1^low^ (ST) SCLC cell lines (mSCLC) cultured in full medium, Cys-depleted medium, or Cys-depleted medium with Fer-1 for 48 h (n=3). **C.** hSCLC cell lines of different subtypes (A, ASCL1^high^; N, NEUROD1 ^high^; P, POU2F3 ^high^) were culture in Cys-depleted medium and stained for lipid ROS accumulation using STY-BODIPY [1 μM] for 48 h (n=3). Levels of lipid peroxidation were monitored by quantifying levels of co-oxidized STY-BODIPY (green, λ_ex_ = 488 nm, λ_em_ = 495−540 nm) emitting in the green spectrum over reduced STY-BODIPY (red, λ_ex_ = 561 nm, λ_em_ = 568−630 nm) emitting in the orange spectrum using the IncuCyte SX5 bioimaging platform. **D.** *RP*-derived mouse ASCL1^high^ (FL) and ASCL1^low^ (ST) SCLC cell lines (mSCLC) were cultured in Cys-depleted medium and stained for lipid peroxidation accumulation using STY-BODIPY [1 μM] for 48 h (n=3). Levels of lipid peroxidation were monitored by quantifying levels of co-oxidized STY-BODIPY (green, λ_ex_ = 488 nm, λ_em_ = 495−540 nm) emitting in the green spectrum over reduced STY-BODIPY (red, λ_ex_ = 561 nm, λ_em_ = 568−630 nm) emitting in the orange spectrum using the IncuCyte SX5 bioimaging platform. **E.** hSCLC cell lines (H82, H526, DMS79) were cultured in Cys-depleted medium, supplemented with Cys, Fer-1 [1 µM], Emricasan (a pan-caspase inhibitor) [2.5 µM], QVD (a pan-caspase inhibitor) [10 µM] and Nec1s (a RIPK1 inhibitor) [10 µM] and stained for lipid ROS accumulation using STY-BODIPY [1 µM] for 48 h (n=3). **F.** Average viability (AlamarBlue assay) for hSCLC cell lines in full medium treated with RSL3 (a ferroptosis inducer) [1 µM] or RSL3 and Fer-1 [1 µM] for 24 h (n=3). **G.** Average cell death (DRAQ7 count/confluence in %) of *RP*-derived mouse ASCL1^high^ (FL) and ASCL1^low^ (ST) SCLC cell lines (mSCLC) cultured in full medium with ferroptosis inducers RSL3 [1 µM], Erastin [10 µM] and ML210 [10 µM] for 24 h, as shown (n=3).

To test whether ASCL1^high^ cells would be resistant to ferroptosis triggered by other means, we used the GPX4 inhibitor RSL3, a *bona fide* inducer of ferroptosis^50^. Again, while human NEUROD1^high^ and mouse ASCL1^low^ SCLC cells were highly sensitive to ferroptosis induced by RSL3, human and mouse ASCL1^high^ cells were generally more resistant to RSL3 treatment (**Figure 4F,G**).

Together, these experiments indicated that subtype-specific sensitivity to ferroptosis can lead to differential dependence on Cys and raised the question how most ASCL1^high^ cells died upon Cys deprivation despite being ferroptosis resistant.

### Cys depletion triggers apoptosis in ASCL1^high^ SCLC cells

We noted that *RPR2*-derived ASCL1^high^ cells showed evidence of caspase 3 cleavage indicative of apoptosis upon Cys deprivation (**Figure 5A**). Indeed, the viability of these ASCL1^high^ cells upon Cys deprivation could not be rescued by Fer-1 (blocking ferroptosis) or by chloroquine (CQ) (blocking autophagy), suggesting that neither ferroptosis nor autophagy are key contributors to cell viability loss in this context; by contrast, a broad-spectrum caspase inhibitor (Boc-D-FMK) as well as GSH supplementation partly rescued viability (**Figure 5B**). *RP*-derived ASCL1^high^ cells also readily induced caspase activity upon Cys depletion (**Figure S7A**) and cell death was partially rescued by co-treatment with the pan-caspase inhibitors emricasan or QVD in these cells (**Figure 4C**). In addition, co-treatment with mitoQ or mitoTEMPO, scavengers for mitochondrial ROS, readily blocked Cys depletion-induced cell death in human ASCL1^high^ cells while it was less effective in NEUROD1^high^ cells (**Figure S7B,C**). Moreover, we observed caspase 3 cleavage in human ASCL1^high^ cells, but not in NEUROD1^high^ cells upon Cys depletion (**Figure 5D**). Notably, cell death induced in human NEUROD1^high^ and POU2F3^high^ cells could not be reverted by caspase inhibition as in human ASCL1^high^ cells (**Figure 5E,F**). Therefore, Cys depletion triggers cell death in ASCL1^high^ SCLC cells at least in part via the induction of mitochondrial ROS and ensuing caspase activation and apoptosis.

**Figure 5:**
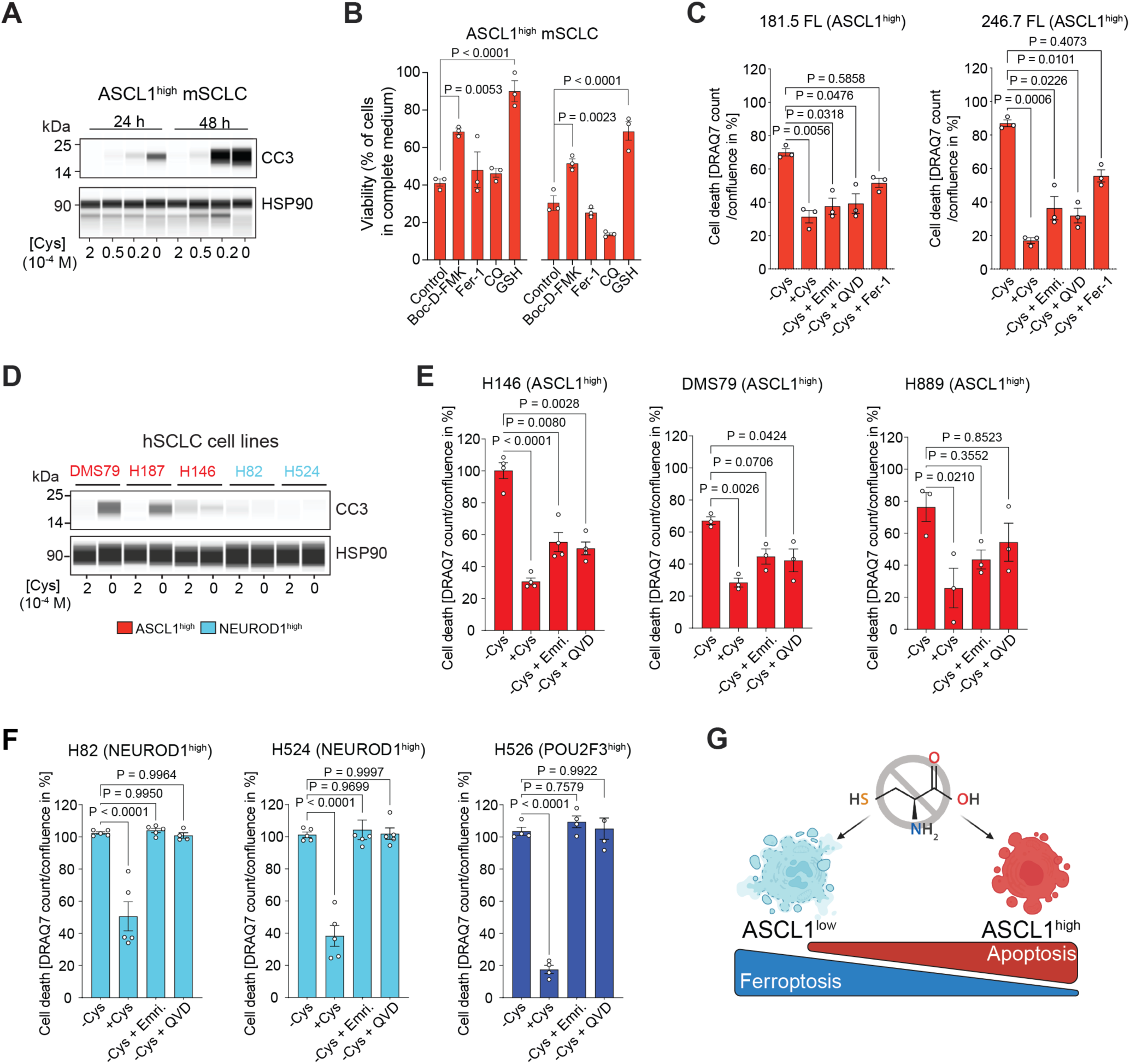
ASCL1^high^ SCLC cells undergo apoptosis in response to Cys deprivation. **A.** Representative immunoassay (n=3) for the marker of apoptosis cleaved caspase 3 (CC3) for ASCL1^high^ mSCLC cells in response to low Cys levels in the medium, as indicated. HSP90 was used as a loading control. **B.** Average viability (AlamarBlue assay) of *RPR2*-derived ASCL1^high^ mSCLC cells cultured in Cys-depleted medium supplemented with Boc-D-FMK [50 µM], Fer-1 [1 µM], chloroquine (CQ) [20 µM], and GSH-MEE [100 µM], as percent of the cells in complete medium (n=3). **C.** Cell death (DRAQ7 count/confluence in %) of *RP*-derived ASCL1^high^ mSCLC cells (181.5 FL, 246.7 FL). Cells were cultured in Cys-depleted medium supplemented with Cys [200 µM], Emricasan (Emri.) [2.5 µM], QVD [10 µM] and Fer-1 [1 µM] for 24 h. **D.** Representative immunoassay (n=3) for CC3 in ASCL1^high^ (NCI-DMS79, NCI-H187, NCI-H146) and NEUROD1^high^ (NCI-H82, NCI-H524) hSCLC cell lines in Cys replete [100 µM] or Cys-depleted medium. HSP90 was used as a loading control. **E,F.** Cell death (DRAQ7 count/confluence in %) of ASCL1^high^ and ASCL1^low^ hSCLC cell lines as indicated. Cells were cultured in Cys-depleted medium, supplemented with Cys [200 µM], Emricasan [2.5 µM], and QVD [10 µM] for 72 h. **G.** Scheme summarizing the consequences of Cys depletion in the different SCLC subtypes. For (B), P-value was calculated by the unpaired Student’s t-test. For (C,E,F) one-way ANOVA was performed, and Tukey was used as post hoc test. Error bars indicate mean ± SEM.

These data reveal that while Cys depletion exploits a metabolic vulnerability in SCLC across cell states, these cell states differ significantly in the type of cell death response triggered with ferroptosis and apoptosis as the dominant forms **(Figure 5G**).

### ASCL1^high^ SCLC cells are ferroptosis resistant due to upregulation of GCH1/BH2/BH4

Given that GSH is needed for GPX4 activity, it came as a surprise that ASCL1^high^ cells, which have low levels of GSH, were resistant to ferroptosis. Intrigued by this observation, we analyzed mRNA expression of known ferroptosis regulators in human ASCL1^high^ cells compared to ASCL1^low^ (NEUROD1^high^ and POU2F3^high^) cells. While the expression of many of these genes was not different between these groups, expression of *GCH1* (coding for GTP cyclohydrolase-1) was elevated in ASCL1^high^ cells (**Figure S8A**). GCH1 is the rate-limiting enzyme for synthesis of 5,6,7,8-tetrahydrobiopterin (BH4), a small metabolite that can act as an endogenous lipophilic radical-trapping agent, thereby protecting from ferroptosis^71,72^ (**Figure 6A**). Expression of *Gch1* was also elevated in the ASCL1^high^ state in matched pairs of RP-derived ASCL1^high^ and ASCL1^low^ SCLC cell lines at the RNA and protein levels (**Figure 6B** and **Figure S8B-C**). Notably, *GCH1* mRNA and protein expression positively correlated with ASCL1 expression across human patient data and human SCLC cell lines **(Figure 6C** and **Figure S8D-G**). In support of elevated GCH1-mediated activity, we noted a significant increase in the levels of 7,8 dihydrobiopterin (BH2), the oxidized form of BH4, in ASCL1^high^ versus ASCL1^low^ mouse SCLC cells (**Figure 6D** and **Figure S8H**), as well as in ASCL1^high^ versus NEUROD1^high^ human cells (**Figure 6E** and **Figure S8I**). Lastly, in a human SCLC scRNA-seq dataset^7^, the highest *GCH1* expression was observed within ASCL1^high^ cells (**Figure S8J**).

**Figure 6:**
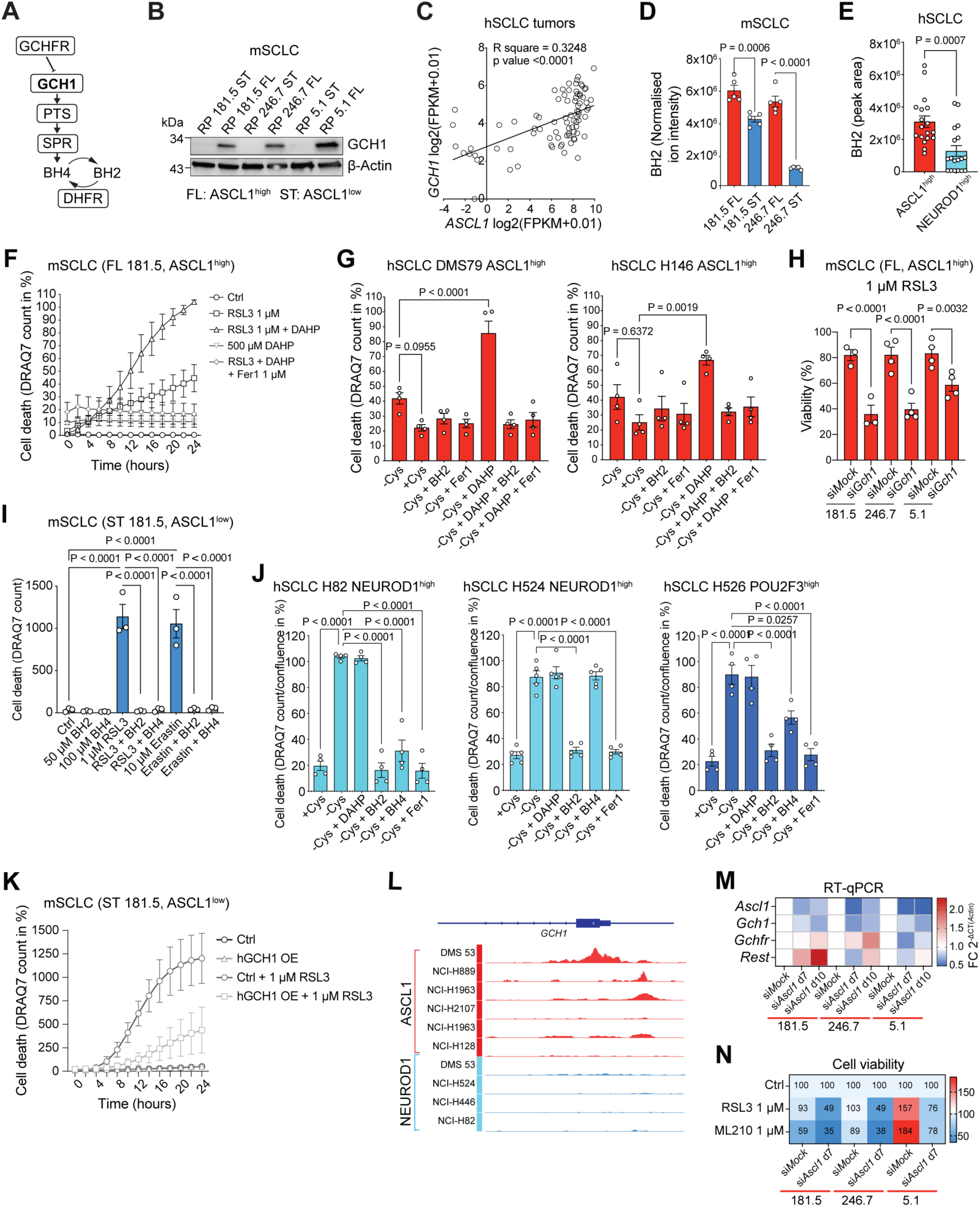
ASCL1^high^ SCLC cells are ferroptosis resistant due to upregulation of GCH1/BH2/BH4. **A.** Schematic representation of the GCH1-BH2/4 metabolic pathway. **B.** Representative immunoassay for in *RP*-derived mouse SCLC cells (181.5 ST/FL, 246.7 ST/FL, 5.1ST/FL). β-Actin was used as a loading control. **C.** Scatter plots of RNA-seq expression data from human SCLC ^44^ for *GCH1* and *ASCL1* expression. **D.** Abundance of BH2 detected by LC/MS in two isogenic pairs of *RP*-derived mouse ASCL1^high^ (FL) and ASCL1^low^ (ST) SCLC cell lines (n=5 technical repeats). **E.** Abundance of BH2 detected by LC/MS in three ASCL1^high^ and three NEUROD1^high^ human cell lines (n=1 experiment per cell line, all 6 technical replicates are shown). **F.** *RP*-derived ASCL1^high^ mouse SCLC cells (181.5 FL) were cultured in full medium or treated with the ferroptosis inducer RSL3 [1 µM], DAHP [500 µM], RSL3+DAHP, and RSL3+DAHP+Fer-1 [1 µM] for 24 h. Cell death (DRAQ7 count in %) was monitored by DRAQ7 [100 nM] incorporation and measured using the IncuCyte SX5 bioimaging platform (n=3). **G.** ASCL1^high^ human SCLC cell lines (hSCLC) were cultured in Cys-depleted medium supplemented with Cys [200 µM], BH2 [50 µM], Fer-1 [1 µM], DAHP [500 µM], DAHP+BH2 or DAHP+Fer-1 for 72 h, n=3. Cell death (DRAQ7 count/confluence) was monitored by DRAQ7 [100 nM] incorporation and measured using the IncuCyte SX5 bioimaging platform (n=4). **H.** *RP*-derived mouse SCLC cells (181.5 FL, 246.7 FL, 5.1 FL) were subjected to GCH1-knockdown (si*Gch1*) or control (siMock) for 5 days were treated with RSL3 [1 µM] for 24 h. Cell viability was measured using CellTiter-Glo (n=3-4). **I.** *RP*-derived mouse SCLC cells (181.5 ST) were treated with RSL3 [1 µM], BH2 [50 µM], BH4 [100 µM], Erastin [10 µM] and the combination of each for 24 h. Cell death (DRAQ7 count) was monitored by DRAQ7 [100 nM] incorporation and measured using the IncuCyte SX5 bioimaging platform (n=3). **J.** hSCLC cell lines (H82, H524, H526) were cultured in Cys-depleted medium, supplemented with Cys [200 µM], BH2 [50 µM], BH4 [100 µM] and Fer-1 [1 µM] for 30 h3. Cell death (DRAQ7 count/confluence in %) was monitored by DRAQ7 [100 nM] incorporation and measured using the IncuCyte SX5 bioimaging platform (n=4-5). **K.** *RP*-derived mouse SCLC cells (181.5 ST) with hGCH1 overexpression were treated with RSL3 [1 µM] for 24 h. Cell death (DRAQ7 count) was monitored by DRAQ7 [100 nM] incorporation and measured using the IncuCyte SX5 bioimaging platform. **L.** Promoter region of human *GCH1* showing published ChIP-seq data of ASCL1 and NEUROD1 (from ^77-80^) binding in different human SCLC cell lines (n=3). **M.** RT-qPCR of *RP*-derived mouse SCLC cells (181.5 FL, 246.7 FL, 5.1 FL) subjected to ASCL1-knockdown (si*Ascl1*) or control (siMock) for 7 or 10 days. Transcripts of *Ascl1*, *Gch1*, *Gchfr* and *Rest1* were measured and plotted normalized to β-actin [FC 2^-ΔCT^] (n=3). **N.** Average viability (CellTiter-Glo) of *RP*-derived mouse SCLC cells (181.5 FL, 246.7 FL, 5.1 FL) subjected to ASCL1-knockdown (si*Ascl1*) or ctrl (siMock) for 7 days were treated with RSL3 [1 µM] or ML210 [10 µM] for 24 h (n=3). For (F,G,H,I,J) one-way ANOVA was performed, Tukey was used as post hoc test. For (E) Student’s t-test was performed. Error bars indicate mean ± SEM.

Next, to determine whether enhanced BH4 synthesis was required for ferroptosis resistance in the ASCL1^high^ state, we treated *RP*-derived ASCL1^high^ cells with RSL3 with and without 2,4-diamino-6-hydroxypyrimidine (DAHP), an activator of GTP cyclohydrolase I feedback regulatory protein (GCHFR), the negative regulator of GCH1 (**Figure 6A**). Strikingly, co-treatment with DAHP was sufficient to sensitize these mouse ASCL1^high^ SCLC cells to RSL3-induced cell death, which was entirely reverted by the ferroptosis inhibitor Fer-1 **(Figure 6F** and **Figure S9A**). Co-treatment with DAHP also sensitized ASCL1^high^ human cell lines to RSL3-induced ferroptosis (**Figure S9B**). Similarly, cell death induced by Cys deprivation in human ASCL1^high^ cells was enhanced by co-treatment with DAHP, and this enhanced cell death was entirely blocked by Fer-1 or BH2 supplementation (**Figure 6G**), indicating a sensitization to ferroptosis induction when BH4 synthesis is inhibited in the ASCL1^high^ state. Treatment with a different BH4 synthesis inhibitor, a sepiapterin reductase inhibitor (SPRi), also sensitized *RP*-derived ASCL1^high^ cells to ferroptosis induction (**Figure S9C,D**). Similarly, siRNA-mediated silencing of GCH1 in mouse ASCL1^high^ cells sensitized these cells to RSL3-induced ferroptosis (**Figure 6H** and **Figure S9E**). Conversely, ferroptosis and lipid ROS induction were abrogated in the presence of BH2 or BH4 in *RP*-derived and human ASCL1^low^ cells both upon RSL3 treatment (**Figure 6I** and **Figure S9F-I**) and upon Cys deprivation (**Figure 6J** and **Figure S9J-L**). Accordingly, overexpression of GCH1 in mouse *RP*-derived and human ASCL1^low^ cells increased cellular BH2 levels and was sufficient to diminish the induction of lipid ROS and render these cells more resistant to ferroptosis (**Figure 6K** and **Figure S10A-G**).

Notably, ASCL1 is crucial for dopaminergic and serotonergic neuronal differentiation^73,74^ and BH4 acts as a cofactor in the synthesis of dopamine serotonin^75,76^. In gene set enrichment analysis (GSEA), we identified that gene sets such as dopamine and serotonin metabolic processes are enriched in ASCL1^high^ versus ASCL1^low^ *RP*-derived SCLC cells (**Figure S10H**). Based on these data, we hypothesized that ASCL1 might directly regulate the expression of GCH1. Indeed, analysis of ChIP-seq data in human SCLC cell lines^77-80^ showed direct binding of ASCL1, but not NEUROD1, to the promoter region of *GCH1* (**Figure 6L**). Accordingly, knockdown of ASCL1 resulted in a decrease of GCH1 RNA and protein levels, and sensitized cells to ferroptosis inducers (**Figure 6M,N**).

These data identify upregulation of the BH2/BH4 synthesis pathway as a mechanism underlying ferroptosis resistance in the ASCL1^high^ state and indicate that upregulation of the ASCL1-GCH1-BH2/4 pathway in SCLC determines whether cells undergo apoptosis or ferroptosis in response to Cys deprivation (**Figure S10I**).

### Cys depletion as the basis for novel treatment combinations in SCLC

Treatment with a cyst(e)inase was previously shown as an effective strategy to induce tumor cell ferroptosis^81^. To further reduce the levels of Cys that may be available to SCLC tumors *in vivo*, we decided to combine injections of PEG-cyst(e)inase with a Cys-depleted chow (“no Cys”). To test the concept of cell-state-dependent ferroptosis sensitization, we treated patient-derived xenograft models (PDXs) from patients with tumors classified as ASCL1^high^, NEUROD1^high^, and POU2F3^high^ ^82,83^. While the NEUROD1^high^ and POU2F3^high^ tumor models showed a moderate response to the no Cys combination treatment, additional treatment with DAHP did not further sensitize these tumors, as predicted from our experiments in cell culture (**Figure7A,B** and **Figure S11A,B**). In contrast, the ASCL1^high^ PDX model did not respond to Cys depletion alone, but showed a clear sensitization in the presence of DAHP (**Figure 7C** and **Figure S11C**). Staining for the lipid peroxidation by-product 4-HNE revealed an increase in signal in the ASCL1^high^ PDX model but not in the two ASCL1^low^ models treated with the combination of Cys depletion and DAHP **(Figure 7D,E** and **Figure S11D-F**). Immunohistochemistry staining for the apoptosis-specific marker cleaved caspase 3 (CC3) confirmed a higher percentage of CC3-positive cells upon Cys depletion in the ASCL1^high^ PDX model, in contrast to the two ASCL1^low^ models (**Figure 7D** and **Figure S11D,G,H**).

**Figure 7:**
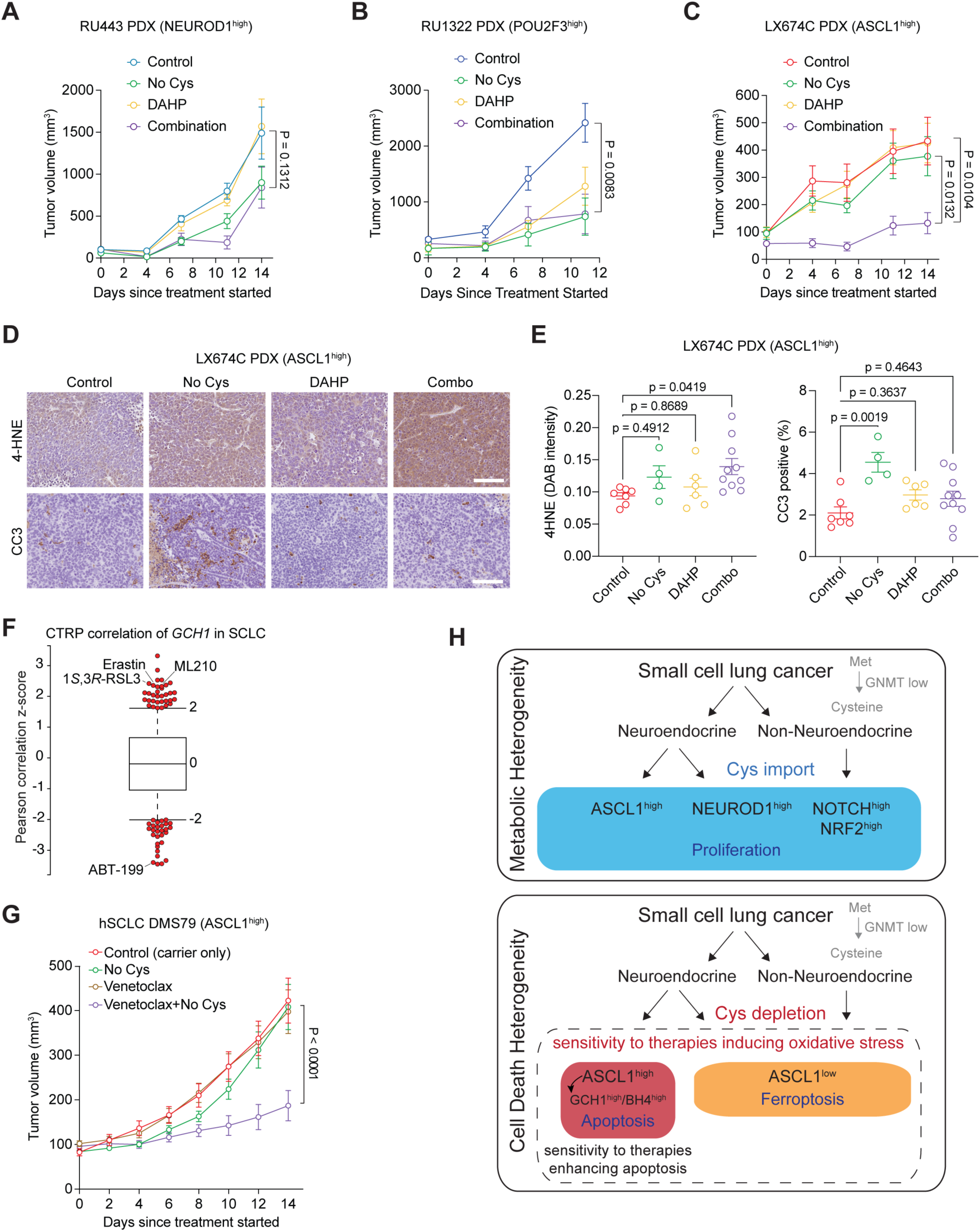
Cys depletion increases treatment susceptibility of SCLC tumors. **A.** Growth of patient-derived xenograft (PDX), LX674C (ASCL1^high^), treated with vehicle, No Cys (cysteinase 50 mg/kg, q.2.d., i.p., and Cys-depleted diet), DAHP (25 mg/kg, q.2.d., i.p.), or both, for 14 days (n=6 mice with two tumors each). **B.** Same as above with PDX, RU443 (NEUROD1^high^). **C.** Same as above with PDX, RU1322 (POU2F3^high^). **D.** Representative images of sections from LX674C tumors immunostained with cleaved caspase 3 (CC3) and 4-HNE antibodies. Scale bars, 100 µm. **E.** Quantifications of 4-HNE and CC3 tumor sections as in (d) (n(control)=7, n(No Cys)=4, n(DAHP)=6, n(combo)=10). **F.** High expression of GCH1 correlates with sensitivity to ABT-199 (venetoclax), and inversely with the ferroptosis inducers ML210, Erastin and RSL3 in human SCLC cell lines. Plotted values are z-scored Pearson’s correlation coefficients; line, median; box, 25th to 75th percentile; whiskers, 2.5th and 97.5th percentile expansion. Data from the CTRP database. **G.** Growth of DMS79 human SCLC xenografts treated with No Cys (cysteinase 50 mg/kg, q.2.d., i.p., and Cys-depleted diet), venetoclax (25 mg/kg, q.i.w., p.o.), or both, for 14 days (n=20). **H.** Model. SCLC tumors are dependent on importing extracellular Cys because they lack expression of the GNMT enzyme in the transmethylation pathway. Under conditions where Cys is available for import, SCLC cells in all states (ASCL1^high^ and NEUROD1^high^ neuroendocrine cells, as well as non-neuroendocrine cells) proliferate and survive. In the absence of exogenous Cys, ASCL1^high^ cells are protected from ferroptosis because ASCL1 directly upregulates the transcription of the gene coding for GCH1, an enzyme critical to produce the antioxidant metabolite BH4. ASCL1^high^ SCLC cells can die from ferroptosis when GCH1 is inhibited or can be pushed towards more apoptotic cell death in the absence of Cys using molecules such as BCL-2 inhibitors. ASCL1^low^ SCLC cells die from ferroptosis upon Cys depletion, and this death can be further enhanced by oxidative stress, including chemotherapy. For (E) one-way analysis of variance (ANOVA) was performed (P<0.0001), and Tukey was used as post hoc test. For all other figures, two-way ANOVA was performed (P<0.0001), followed by the unpaired Student’s t-test. Error bars indicate mean ± SEM.

While these data supported the concept of exploiting Cys depletion for the treatment of SCLC, DAHP does not currently meet requirements for clinical testing. To find combinatorial strategies specifically tailored towards SCLC tumors that are more resistant to ferroptosis (i.e., tumors with ASCL1^high^ and GCH1^high^ cells), we mined the Cancer Therapeutics Response Portal (CTRP)^84-86^. This analysis confirmed that high expression of *GCH1* in human SCLC cell lines correlated with resistance towards ferroptosis-inducing small molecules (e.g., RSL3, ML210, Erastin) (**Figure 7F**) and identified the BCL-2 inhibitor and intrinsic apoptosis inducer venetoclax (ABT-199) as a top candidate correlating with sensitivity (**Figure 7F**). Conversely, the efficacy of ABT-199 treatment response in all the human SCLC cell lines in this database correlated with high *GCH1* expression (**Figure S12A**). ASCL1 expression was previously shown to correlate with BCL-2 expression and ABT199 response^87,88^, and we found that resistance to Cys depletion-induced apoptosis correlated with BCL-2 expression in ASCL1^high^ cells (**Figure S12B**).

Based on these observations, we tested the combination of Cys depletion and apoptosis induction upon BCL-2 inhibition. We found that Cys depletion increased the sensitivity of ASCL1^high^ human SCLC cell lines to venetoclax in culture (**Figure S12C**). In DMS79 xenografts, the regimen we used for venetoclax monotherapy did not change tumor growth, but the combinatorial strategy with Cys depletion significantly reduced tumor growth, without body mass loss (**Figure 7G,H** and **Figure S12F-H**). The anti-tumor effects of the combination treatment were accompanied by increased apoptosis (**Figure S12I,J**). Thus, promoting apoptosis through venetoclax treatment and Cys deprivation in ferroptosis-resistant ASCL1^high^ SCLC cells can limit the growth of SCLC tumors.

Overall, our data reveal that Cys depletion can be effectively combined with other strategies to inhibit tumor growth in pre-clinical models of SCLC (**Figure 7H**).

## DISCUSSION

A single-cell transcriptional analysis across 24 tumor types identified the glutathione pathway as a core “meta-program” related to intratumoral heterogeneity ^36^, but the relevance of this observation for tumor progression and therapy has remained unclear. Through metabolic pathway mapping, we illustrate the differential regulation of the Cys-glutathione axis between cancer cell states defined by ASCL1 expression in SCLC, identifying cancer cell populations with varying redox conditions and Cys requirements for survival. This observation underlines heterogenous levels of oxidative stress within a tumor may affect responses to conventional therapies that rely on oxidative stress induction, including radiation therapy and chemotherapy.

The *de novo* synthesis pathway can normally produce cysteine in cells, but the essentiality of extracellular Cys for cancer cell proliferation has long been recognized^89,90^. Our data further reveal that the dependency on extracellular Cys in cancer cells stems from insufficiency in endogenous cysteine synthesis. Interestingly, GNMT is a tumor suppressor in liver cancer through distinct pathways^91^. Future experiments may seek to better understand why SCLC cells do not evolve to express higher levels of GNMT, especially upon radiation therapy and chemotherapy treatment, as low levels of this enzyme may increase cancer cell death under conditions of oxidative stress.

While the induction of ferroptosis is viewed as a promising strategy to inhibit tumor growth, the best way to implement this idea in cancer remain unclear, in part because the endogenous mechanisms used to manage oxidative stress in tumors are still not certain^92-94^. Recent studies have shown that oncogenic transcription factors can drive ferroptosis sensitivity^60,95^, as do the pools of cysteine in distinct intracellular compartments^96^. Interestingly, we find that elevated expression of GCH1 and the synthesis of its product BH4 cause ferroptosis resistance in the ASCL1^high^ state in SCLC. The process of ASCL1-induced transcription of GCH1 can be thought of as a “neuroendocrine heritage” of SCLC, as BH4 is involved in serotonin and dopamine biosynthesis, two important neuroendocrine transmitters^76^. Thus, ferroptosis resistance in ASCL1^high^ SCLC might be an evolutionary by-product passed down from the cell-of-origin or related to the neuroendocrine state of these cancer cells. ASCL1-driven neuronal differentiation was shown to pass through a ferroptosis checkpoint, wherein overcoming ferroptosis helped to obtain more differentiated ASCL1-expressing neurons^97^. Taking these data into consideration, GCH1 expression might also be required for ASCL1^high^ SCLC cells to withstand a putative bottleneck of lipid oxidative stress during NE differentiation and plasticity.

Other studies have explored subtype-specific susceptibilities to purine synthesis and glycolysis inhibitors^98,99^, as well as arginine depletion^100^ in SCLC. Moreover, ASCL1^high^ SCLC was recently shown to depend on oxidative phosphorylation (OXPHOS)^101,102^. Cys dependency of ASCL1^high^ cells might therefore be a means to cope with elevated levels of ROS coming from OXPHOS. It is likely that efficient therapeutic approaches against SCLC will need to inhibit the expansion of most, if not all, cell states, and our data suggest that Cys depletion, which inhibits the expansion of all the cell states we examined, could prove a worthwhile strategy to explore clinically. However, the ability of the liver to synthesize Cys that can be used by other organs and, potentially, tumors^67^, may limit the ability of body-wide Cys depletion to inhibit tumor growth. To address this issue at least partly, we implemented a combined strategy involving dietary and enzymatic depletion of Cys^65^. Other strategies may include targeting the x_c_^-^ cystine/glutamate antiporter^103^. We found that Cys depletion alone following our protocol only moderately inhibited tumor growth. To achieve better control of tumor growth, combination of Cys depletion and another therapeutic molecule such as cisplatin or venetoclax was necessary. Depletion of cysteine or other amino acids may help treat SCLC^104,105^ ^22^ and other cancer types^106-108^ ^62,109-111^. However, it is likely that in all, these cases, combinatorial approaches with additional treatments will be required for consistent therapeutic outcomes. Considering that modulating oxidative stress is a crucial mechanism underlying many cancer treatments and drug resistance^45^ ^112^, we still anticipate that inhibition of Cys and the associated pathways in cancer holds potential as an adjuvant therapy for diverse treatments, allowing both broad and tailored strategies for different cancer types.

## METHODS

### Ethics statement

Mice were maintained according to practices approved by the NIH, the Stanford Institutional Animal Care and Use Committee (IACUC), and the Association for Assessment and Accreditation of Laboratory Animal Care (AAALAC). The study protocol was approved by the Stanford Administrative Panel on Laboratory Animal Care (APLAC) (protocol 13565).

### SCLC development in mutant mice

Tumor initiation and development in *Rb^flox/flox^*;*p53^flox/flox^*;*Rbl2^flox/^*^flox^;*Hes1^GFP^*mice following Ad-Cre delivery (4x10^7^ PFU of Adeno-CMV-Cre, Baylor College of Medicine, Houston, TX) has been described previously ^16^. Mice were euthanized before or at the first sign of respiratory distress in accordance with Stanford APLAC guidelines. Mice were housed at 22 °C ambient temperature with 40% humidity and a light/dark cycle of 12 h (7am-7pm).

### Flow cytometry sorting of SCLC cells

Cells were isolated from microdissected *RPR2* mutant tumors as described^16^. Briefly, pooled tumors were minced with a razor blade, digested with collagenase/dispase (Roche) and then cooled on ice with DNase1 (Millipore) before cells were passed through a 40 µm filter, spun down, and resuspended in 1 mL of red blood cell lysis buffer. Cells were then resuspended in 10% BGS (Fisher Scientific, Bovine Growth Serum) in Phosphate Buffered Saline (PBS), at 10^6^ cells per 100 µL) and stained. We used CD45-PE-Cy7 (eBioscience, 25-0451-82, clone 30-F11, 1:100), CD31-PE-Cy7 (eBioscience, 25-0311-82, clone 390, 1:100), TER-119-PE-Cy7 (eBioscience, 25-5921-82, clone TER-119, 1:100), CD24-APC (eBioscience, 17-0242-82, clone M1/69, 1:200) or CD45-Pacific Blue (BioLegend, 103126, clone 30-F11, 1:100), CD31-Pacific Blue (BioLegend, 102422, clone 390, 1:100), TER-119-Pacific Blue (BioLegend, 116232, clone TER-119, 1:100), CD24-PE-Cy7 (BioLegend, 101821, clone M1/69, 1:200), and DAPI to label dead cells. Cells were sorted with a 100 µm nozzle on a BD FACSAria II using the FACSDiva software. Data were analyzed and plotted using FlowJo v10.

### Cell culture

All cell lines (Human SCLC cell lines: H82, H524, H526, H1694, H446, H2227, H2018, DMS79, H187, H146, H889; *RP* mutant mouse SCLC cell lines: 181.5 ST/FL, 246.7 ST/FL, 5.1 ST/FL (as in ^21^); KP1 ^113^; *RPR2* mutant cell lines: HES1-GFP^pos^ and HES1-GFP^neg^ cell lines from three independent tumors that we internally labelled as #326, #330, and #1157) were grown in RPMI-1640 medium supplemented with 10% bovine growth serum (BGS) (Fisher Scientific, SH3054103HI) and (Fisher Scientific, 10378016). For Cys-depleted conditions, RPMI without methionine, cystine and l-glutamine (Millipore-Sigma, R7513) was used. For Cys-depletion experiments, the medium was supplemented with and 2% BGS and 1% penicillin-streptomycin-glutamine and 100 µM methionine and 2 mM l-glutamine as in full RPMI-1640 media.

The following molecules were used in cells: RSL3 (Cayman Chemical, 19288), ML210 (Tocris Bioscience, 6429/10), Erastin (Cayman Chemical, cay17754-5), venetoclax (Selleck Chemicals 8048), cisplatin (Mount Sinai Hospital Pharmacy), Emricasan (Cayman Chemical 12986), z-VAD(OMe)-FMK (MedChemExpress, HY-16658), Nec1s (Abcam, ab221984), propidium iodide (Millipore-Sigma, P4170), QVD (MedChemExpress, HY-12305), GSH-MEE (Millipore-Sigma, G1404), SPRi (MedChemExpress, HY-115510), DAPI (Millipore-Sigma, D9542), DRAQ7 (BioLegends, 424001), ferrostatin-1 (Yaman Chemical, 17729-25), a-tocopherol (Cayman Chemical, 25985), ethacrynic acid (Cayman Chemical 19536), ascorbic acid (Spectrum S1349), methionine (Millipore-Sigma, M5308), sulfasalazine (Millipore-Sigma, S0883), STY-BODIPY (Cayman Chemical, Cay27089-500), puromycin (Thermo Fisher Scientific, A11138), ampicillin (Millipore-Sigma, A0166-5G) and blasticidin (Thermo Fisher Scientific, R21001).

### Knock-down and overexpression studies in cells

shRNAs were obtained from the MISSION shRNA library (Millipore-Sigma). Scramble shRNA (CAACAAGATGAAGAGCACCAA) cloned into the backbone pLKO was used as a control. For *Nrf2/Nfe2l2*, the sequences of the shRNAs are 1: 5’-GCAGAATTATAGCCAAAGCTA and 2: 5’-CCACGCTGAAAGTTCAGTCTT. The lentiviral vector for GNMT expression was purchased from DNASU (HsCD00433897). siRNA mediated knockdowns were performed using the Dharmacon siRNA system. For this, 200 μL Opti-MEM were mixed with 1.5 μL Dharmafect reagent I and incubated for 5 min at RT. 2.8 μL of the respective 20 μM siRNA were added and incubated for another 30 min. 200 μL siRNA mix was added in a well of a 6-well plate first, and cells were added in 1 mL antibiotic free medium on top. Knockdowns were incubated for 48–72 h, as indicated. If gene silencing was carried out over several days, a fresh siRNA mixture was added every second day. ON-TARGETplus Smartpool siRNAs (mouse *Ascl1* siRNAs: 5’-CAACGUCAGUUCUCGGACA-3’; 5’-ACAGAAACGUGGUUAAUGU-3’; 5’-ACAGACAGACACUAUAUUA-3’; 5’-CCGAAUCACAGAUGGGUUC-3’); mouse *Gch1* siRNA (5’-GGUAGAAUGCUAAGUACGU-3’; 5’-CGAGAAGUGUGGCUACGUA-3’; 5’-GAGAAGGGAGAAUCGCUUU-3’; 5’-AGUAGUGAUUGAAGCGACA-3’), mouse mock siRNA 5’-UGGUUUACAUGUCGACUAA-3’; 5’-UGGUUUACAUGUUGU GUGA-3’; 5’-UGGUUUACAGUUUUCUGA-3’; 5’-UGGUUUACAUGUUUUCCUA-3’) were obtained from

Dharmacon.

### Preparation and isolation of plasmid DNA

For isolation of plasmid DNA from *Escherichia coli* (*E. coli*), bacteria were cultured for approximately 7 h in a shaking incubator at 37°C and 1500 x g in 5 mL lysogeny broth (LB). This culture was used to inoculate 200 mL LB medium supplemented with 50 μg/mL ampicillin which was grown overnight. Plasmid DNA was isolated using E.Z.N.A.® Plasmid Maxi Kit according to the manufactureŕs instructions.

### Transfection of plasmid DNA

One day before transfection, cells were seeded in 2 mL of the respective medium without antibiotics so that cells would be 70-90% confluent at the time of transfection. For transfection of plasmid DNA, 250 μL Opti-MEM were mixed with 10 μL Lipofectamine 2000 per well of a 6-well plate. In an additional tube 4 μg plasmid DNA (h*GCH1* expression vector was purchased from VectorBuilder) were diluted in 250 μL Opti-MEM and incubated for 5 min at RT. Subsequently, diluted DNA was added to the Opti-MEM Lipofectamine mix and incubated for 20 min at RT. Then, 500 μL of the transfection mix was dropped on the cells. After six hours, the medium was replaced with fresh medium. Cells were tested for transgene expression after 24 h and 48 h.

### Chromatin immunoprecipitation (ChIP)

ChIP was performed as previously described^3^. Briefly, doxycycline (0.125 µg/mL) induced KP1-pLIX-N1ICD cells were fixed with 2 mM disuccinimidyl glutarate (Thermo Fisher Scientific) in PBS and formaldehyde. The antibodies used were Notch1 (CST, 3608) and rabbit IgG (CST, 2729). For detection of mouse Nrf2 promoter, DNA solution was used as a template with the following primers: 5’-GAGCGACCTTCGGCAAACA-3’ and 5’-GGGCATGGACCTGAGTGCTT-3’.

### Quantitative real-time PCR (RT-qPCR)

For the data presented in Figure 1, RNA was isolated using the RNeasy mini kit (Qiagen 74106) according to the manufacturer’s protocol. RT-qPCR analysis was performed on the Applied BiosystemsTM 7900HT Fast Real-Time PCR system using PerfeCTa SYBR Green FastMix (Quanta BioSciences, 95073). Mouse *Nrf2* expression was detected using the following primers: 5’-CTTTAGTCAGCGACAGAAGGAC-3’ and 5’-AGGCATCTTGTTTGGGAATGTG-3’. Data was normalized to *Gapdh* as a house keeping gene using the following primers 5’-AGGTCGGTGTGAACGGATTTG-3’ and 5’-GGGGTCGTTGATGGCAACA-3’.

For the data presented in Figure 6, the NucleoSpin RNA kit (Macherey-Nagel, 740955.5) was used according to the manufacturer’s protocol. The isolated RNA was reverse transcribed into cDNA using the LunaScript RT SuperMix Kit (NEB, E3010L) following the protocol provided by the manufacturer. For quantitative PCR the Power SYBR Green PCR Master Mix (Thermo Fisher Scientific, 4368702) was used. To this end, 5 μL Power SYBR Green PCR Master Mix was mixed with 2 μL of nuclease-free water (NEB). After adding 1 μL (10 μM) of forward and reverse primers (5’>3’: mβ-Actin fw: GGCTGTATTCCCCTCCATCG, mβ-Actin rv: CCAGTTGGTAACAATGCCATGT; mAscl1 fw: GCAACCGGGTCAAGTTGGT, mAscl1 rv: CAAGTCGTTGGAGTAGTTGGG, mGch1 fw: GCCGCTTACTCGTCCATTCT, mGch1 rv: GAACAAGGTGATGCTCACACA; mGchfr fw: GCACTCAGATCCGTATGGAGG, mGchfr rv: GTTGTTTCCCAAGACACTTCTCT; mRest fw: AGCGAGTACCACTGGAGGAA, mRest rv: CTGAATGAGTCCGCATGTGT) and 2 μL of cDNA (5 μg/μL) real-time qPCR was performed in triplicates on the Quant Studio5 RT-qPCR machine. Results were normalized to the expression of the house-keeping gene Actin.

### Immunoblotting

Cells were washed, lysed in RIPA-lysis buffer (Thermo Fisher Scientific, 89901) plus Protease and Phosphatase inhibitor (Roche) and frozen at -20°C. After re-thawing, samples were centrifuged at 18,000 x g for 20 min at 4°C. Lysate concentrations were adjusted to equal protein concentrations using the bicinchoninic acid (BCA) protein assay (Bio-Rad). Equal amounts of protein were mixed to a final concentration of 1x reducing sample buffer (Invitrogen) containing 200 mM DTT. Samples were heated to 80°C for 10 min, separated via gel electrophoresis and transferred to Nitrocellulose membranes using the TurboBlotting system (Bio-Rad). Membranes were blocked in PBS with 0.1% Tween 20 (PBST) with 5% (w/v) bovine serum albumin (BSA) for at least 30 min. Next, membranes were incubated over night at 4°C with primary antibodies (Anti-ß-actin (Millipore-Sigma, A1978, 1:10,000), Anti-ASCL1 (BD Pharmingen, 556604, 1:1,000), Anti-REST (Thermo Fisher Scientific, BS-2590R, 1:1,000), Anti-YAP1 (Cell Signaling Technology, #4912, 1:1,000), Anti-NEUROD1 (Abcam, ab60704, 1:1,000), Anti-hGCH1 (Abcam, ab236387, 1:1,000), Anti-mGCH1 (Abcam, ab307507, 1:1,000), Anti-GCHFR (Thermo Fisher Scientific, 18809-1-AP, 1:1,000)) in PBST with 5% BSA. After washing with PBST, membranes were incubated with horse radish peroxidase (HRP)-coupled secondary antibodies for at least 1 h. After another washing step, membranes were developed using chemiluminescent Amersham ECL Prime Western Blotting Detection Reagent (Cytiva, RPN2235).

### Recombinant cyst(e)inase

Mouse, human, and engineered cystathionine gamma lyases containing N-terminal hexahistidine tag was expressed as previously described^65^. Briefly, cDNA encoding for mouse, human, and engineered cystathionine gamma lyase (CGL) without the signaling peptide sequence was cloned into pBAD plasmid with an inducible araBAD promoter using *EcoR*I and *Xho*I restriction sites and amplified in E. coli-BL21(DE3) cells. Cells in the logarithmic phase of growth were induced with 0.1% (w/v) arabinose in Terrific Broth (Thermo Fisher Scientific, 22711022) for 10 h at 37°C. Induced cells were pelleted and resuspended in B-PER reagent (Thermo Fisher Scientific, 90084) with lysozyme (0.1 mg/mL) and DNase I (5 U/mL) and incubated for 10 min at 37°C. Cyst(e)inases were then purified using a nickel-NTA affinity column (Qiagen, 30210) and dialyzed into PBS (Thermo Fisher Scientific, 66110). PEGylation was conducted using 100-fold molar excess of methoxy PEG succinimidyl carboxymethyl ester, MW 5kDa (JenKem Technology, A3012-1).

### Subcutaneous tumor studies

For allograft studies, B6129SF1/J immunocompetent mice (The Jackson Laboratory, Strain #:101043) were implanted with 1 x 10^6^ mouse SCLC cells in 50% Matrigel into the lower left and right quadrants. The tumors were allowed to reach 100 mm^3^. The mice were then randomized according to the tumor size and treated (50 mg/kg q.2.d., i.p.) with the functional engineered cyst(e)inase or the boiled enzyme (90°C, 5 min) for 14 days. For combinatorial treatment with cisplatin (Mount Sinai Hospital Pharmacy), allografts were treated (4 mg/kg, q.wk., i.p.) with or without the same dosing regimen of the engineered cyst(e)inase. Dietary depletion of glutamine or Cys was initiated at the time of tumor injection by transitioning to 0.0% w/w glutamine or 0.0% w/w Cys chows (Research Diets).

For xenograft studies Nod.Cg-Prkdc^scid^IL2rg^tm1Wjl^/SzJ (NSG) mice were implanted with 1 x 10^6^ cells in 50% Matrigel and allowed to reach 100 mm^3^ before randomization. Mice with NCI-H82 xenografts were treated with Cys depletion (50 mg/kg cyst(e)inase q.2.d., i.p. and Cys-depleted chow), cisplatin (4 mg/kg, b.i.w., i.p.), or Cys depletion and cisplatin combined. For combinatorial treatment with venetoclax (Selleck Chemicals, 8048), mice with NCI-DMS79 xenografts were treated with Cys depletion (50 mg/kg cyst(e)inase q.2.d., i.p. and Cys depleted chow), venetoclax (25 mg/kg, q.i.w., p.o.), or Cys depletion and venetoclax combined. Both male and female mice were used without selection for sex.

For combinatorial treatment with DAHP (MedChemExpress, HY-100954), we used patient-derived xenografts (PDXs) (MSK443, MSK674c, and MSK1322 from the Rudin Lab, IACUC-approved protocol 04-03-009). MSK443 and MSK674c were published in ^83^, and MSK1322 was leveraged in ^82^. Tumors were allowed to grow until they reached 100-200 mm^3^, at which point, they were randomized into groups and treated with either vehicle, no Cys (50 mg/kg q.2.d., i.p. and Cys-depleted diet), DAHP (25 mg/kg, q.2.d., i.p.), or a combination of no Cys and DAHP.

### Metabolite extraction and LC-MS analysis of ASCL1^high^ and ASCL1^low^ mouse SCLC cells

SCLC cells sorted using flow cytometry (see Flow cytometry sorting of SCLC cells), cells were expanded in RPMI-1640 medium supplemented with 10% BGS (see Cell culture). For each condition, the cells were treated in 6-well plates, centrifuged at 300 rcf for 5 min at 4°C and washed twice with ice-cold PBS. Metabolites were extracted by immediately adding 300 µL of precooled 80% MeOH/H_2_O to cell pellets. After keeping the cells in -80°C for 15 min, they were vortexed and centrifuged at 20,000 rcf for 10 min at 4°C. The supernatant was collected for LC-MS analysis.

LC/MS analyses were performed using positive and negative ion mode respectively on an Agilent 1290 Infinity LC system coupled to an Agilent 6545 Q-TOF mass spectrometer with an Agilent Jet Stream Source. The LC chromatographic separation was optimized to achieve the optimal selectivity on an Agilent InfinityLab Poroshell 120 HILIC-Z (2.1 mm × 150 mm, 2.7 µm) PEEK-lined column. In brief, for the positive ion mode, the mobile phase A was 10 mM ammonium formate in water with 0.1% formic acid, and the mobile phase B was 10 mM ammonium formate in acetonitrile/water (90:10 v/v) with 0.1% formic acid. The gradient elution was 0-3 min, 98% B; 3-11 min, 98% to 70% B; 11-12 min, 70%-60% B; 12-14 min, 60%B; 14-14.1min, 60%-98% B; 14.1-16 min, 98% B. For the negative ion mode, the mobile phase A was 10 mM ammonium acetate in water with 2.5 µM deactivator additive (pH=9) and the mobile phase B was 10 mM ammonium acetate in acetonitrile/water (85:15 v/v) with 2.5 µM deactivator additive (pH=9). The gradient elution was 0-2 min, 96% B; 2-5.5 min, 96%-88% B, 5.5-8.5 min, 88% B; 8.5-9 min, 88%-86% B; 9-14 min, 86%B; 14-17min, 86%-82% B, 17-23 min, 82%-65% B; 23-24 min, 65% B; 24-24.5 min, 65%-96% B; 24.5-26 min, 96% B; post time was 3 min. Other key LC parameters were: LC flow rate was 0.25 mL/min; autosampler temperature was 4°C; injection volume was 3 µL and column temperature was 25°C for positive and 50°C for negative ion mode. The 6545 Q-TOF mass spectrometer parameters were optimized to achieve the best sensitivity and a broad coverage of metabolites. The mass spectrometer parameters were as follows: acquisition mass range was 60-1200 m/z, Vcap was 3000 V for the positive ion mode and 3500 V for negative ion mode, gas temperature was 225°C at a flow of 13 L/min and Sheath gas temperature was 225°C for the positive ion mode and 350°C for the negative ion mode with flow of 12 L/min, nebulizer is 35 psi, fragmentor is 125 V. Dynamic mass axis correction was achieved by continuous infusion of a reference mass solution including two ions at m/z= 121.05087 and m/z=922.0098 for the positive ion mode and m/z= 68.9957 and m/z=980.0164 for the negative ion mode.

The following molecules were used as standards: L-cysteine (Millipore-Sigma, C7352), L-cystine (Millipore-Sigma, C8755), L-glutathione (Millipore-Sigma, G4251), oxidized L-glutathione (Millipore-Sigma, G4376), L-homocysteine (Millipore-Sigma, 69453), s-adenosylhomocysteine (Millipore-Sigma, A9384), s-adenosyl methionine (S Millipore-Sigma, 69453), gamma glutamyl cysteine (Millipore-Sigma, G0903), L-cysteine sulfinic acid (Millipore-Sigma, C4418), hypotaurine (Millipore-Sigma, H1384), taurine (Millipore-Sigma, T0625) and dihydrobiopterin (Cayman 81882).

### Metabolomics data analysis of mouse *RPR2* mutant SCLC cells

For the metabolomics data analysis, Agilent MassHunter Profinder software B.08.00SP3 was used for batch processing of the accurate mass Q-TOF LC-MS data. In this study, Agilent Pathways to Personal Compound Database and Library (PCDL) software B.07.00 was used to create an Agilent PCDL file (.cdb) from Wiki and BioCyc metabolic pathways. The created PCDL file was then edited in Agilent PCDL Manager software B.08.00 to remove and add some metabolites based on the study needs. The final database containing 2966 metabolites was used as a target list for Batch Targeted Feature Extraction in MassHunter Profinder software. After batch targeted feature extraction, the results were reviewed and manually curated as necessary. The final curated results with 1296 metabolites were exported as a .pfa file that was then imported into Agilent Mass Profiler Professional (MPP) software 15.1 for statistical analyses such as PCA, Volcano plot and Hierarchical clustering. The pathway analysis was performed for those statistically significant metabolites with P≤0.005 and fold change (FC) ≥2 using Pathway Architect included in the MPP software.

Metabolites were separated chromatographically using a Millipore SeQuant ZIC-pHILIC analytical column (5 µm, 2.1 × 150 mm) with a matching 2.1 × 20 mm guard column, both with 5 µm particle size. A binary solvent system was employed, consisting of 20 mM ammonium carbonate with 0.05% ammonium hydroxide as Solvent A and pure acetonitrile as Solvent B. Chromatography was performed at 40°C with a flow rate of 0.200 mL/min, using the following gradient program: 80% Solvent B from 0-2 min, a linear gradient to 20% Solvent B from 2-17 min, a rapid return to 80% Solvent B from 17-17.1 min, and a hold at 80% Solvent B from 17.1-23 min. The autosampler tray was maintained at 4°C, and 5 µL of each sample was injected. Samples were randomized to minimize analytical variability, and pooled quality control (QC) samples, prepared by mixing equal volumes of all individual samples, were analyzed regularly to ensure consistency.

### Metabolomics data analysis of mouse *RP* mutant SCLC cells

Metabolite quantification was performed using a Vanquish Horizon UHPLC system coupled to an Orbitrap Exploris 240 mass spectrometer (Thermo Fisher Scientific) with a heated electrospray ionization (HESI) source. Ionization parameters were set to +3.5 kV (positive mode) and -2.8 kV (negative mode), with an RF lens value of 70, a heated capillary temperature of 320 °C, and an auxiliary gas heater temperature of 280 °C. Sheath gas flow was set to 40, auxiliary gas to 15, and sweep gas was off. MS1 scans covered a mass range of m/z 70-900, with automatic gain control (AGC) targets and maximum injection times. Data were acquired in full scan mode with polarity switching at a resolution of 120,000. The AcquireX Deep Scan workflow facilitated untargeted metabolite identification via iterative data-dependent acquisition. Fragmentation was conducted with resolutions of 60,000 for full scans and 30,000 for MS/MS, an intensity threshold of 5.0 × 10^3^, a 10-second dynamic exclusion, a 5 ppm mass tolerance, an isolation window of 1.2 m/z, and stepped normalized collision energies of 30, 50, and 150. Mild trapping was enabled to improve signal quality.

Metabolite identification was performed using Compound Discoverer software (v3.2, Thermo Fisher Scientific), with criteria including a precursor ion m/z within 5 ppm of the theoretical mass, fragment ions within 5 ppm of an internal spectral library, a minimum match score of 70, and retention times within 5% of purified standards analyzed under identical conditions. Peak area integration and chromatogram review were carried out using TraceFinder software (v5.0, Thermo Fisher Scientific).

Peak areas were normalized against the total ion count (TIC) of each sample to correct for analytical variability, and the resulting normalized values were used for statistical analyses. Principal component analysis (PCA) was performed using the prcomp function from R’s stats package (https://www.R-project.org/), and plots were visualized with ggplot2^114^. Sample group differential abundance was assessed using two-sided Student’s t-tests, and false discovery rate (FDR) correction was applied using the Benjamini–Hochberg method from R’s base stats package. EnhancedVolcano^115^ was used to generate volcano plots for visualization of the differential abundance analysis. MetaboAnalyst 4.0^116^ was used for over-representation analysis (ORA). Boxplots were created using ggstatsplot^117^. Heatmaps were generated with the pheatmap package^118^. Finally, tidyverse was used for general scripting^119^.

### Metabolic tracing of human SCLC cells

RPMI medium containing [3-^13^C] serine (Cambridge Isotope, CLM-1572) was prepared following the standard RPMI formula (US Biological Life Sciences, R8999-04A) without serine and supplemented with isotopic serine or unlabeled serine and 10% dialyzed FBS. For Cys depleted medium, the same formula was used but cystine was excluded. Cells were cultured in 6-well plates with the isotopic medium for the indicated time, then washed with ice-cold PBS twice, and metabolites were extracted by immediately adding 300 µL of 80% acetonitrile/H_2_O. The samples were kept on ice for additional 15 min. The cell suspension was vortexed and centrifuged at 20,000 rcf for 10 min and stored at -80C prior to LC-MS analysis.

For the metabolic tracing study involving GNMT overexpression, LC-MS analysis was conducted as above. An Agilent PCDL file (.cdb) was created for a list of interesting metabolites. This PCDL file was edited by adding retention times for each metabolite. The PCDL file was then used as a target list (**Table S4**) for batch isotopologue extraction in Agilent MassHunter Profinder software. The batch isotopologues results were corrected for 13C natural abundance and the [3-^13^C] Serine isotopic purity (99%).

For the metabolic tracing study involving homocysteine, 200 µL of supernatants were subjected to LC-MS analysis. Liquid chromatography was performed using an Agilent 1290 Infinity LC system coupled to a 6546 Q-TOF mass spectrometer. A hydrophilic interaction chromatography method with a HILIC column (100 x 2.1 mm, 3 µm; Waters) was used for compound separation with column temperature at 35°C and a flow rate of 0.3 mL/min. Mobile phase A consisted of 10 mM ammonium formate and 0.1% formic acid in water and mobile phase B was 10 mM ammonium formate and 0.1% formic acid in water:acetonitrile (1:9). The gradient elution was 0—1 min, 85% B; 1—9 min, 85% B → 65% B, 9—13 min, 65% B → 35% B, 13—16.5 min, 35% B.

The overall runtime was 17 min, and the injection volume was 5 µL. The Agilent Q-TOF was operated in positive ion mode, and the relevant parameters were as listed: Vcap, 3500 V; nozzle voltage, 1000 V; fragmentor voltage, 125 V; drying gas flow (350°C), 11 L/min; sheath gas temperature, 325 °C; and nebulizer pressure, 40 psi. Acquisition mass range was 50-1600 (m/z). The reference masses were 121.0509 and 922.0098. The acquisition rate was 2 spectra/s. Targeted analysis, isotopologues extraction (for the metabolic tracing study), and natural isotope abundance correction were performed by the Agilent Mass Hunter Profinder B.10.00 Software.

### Desorption electrospray ionization (DESI) mass spectrometry imaging (MSI) analysis of *RPR2* SCLC mouse model

After tumor development in *RPR2* mutant mice following Ad-Cre delivery, lung specimens were collected for immediate freezing in O.C.T. (Sakura TissueTek, O.C.T. Compound 4583) with liquid nitrogen into cryoblocks that were kept at -80 °C until sectioning. The cryoblocks were sectioned at 10 µm thickness. Due to the large ion suppression by O.C.T., analysis was performed using negative ionization mode. DESI MSI data were obtained from vacuum desiccated tissue sections placed on a DESI 2D^TM^ XS (Waters Corporation) coupled with a high-resolution quadrupole time-of-flight mass spectrometer (SELECT SERIES^TM^ Cyclic IMS mass spectrometer, Waters Corporation). DESI data was acquired using 98% methanol and 2% aqueous solvent with 0.1% *v/v* formic acid and 200 pg/µL of Leu-enkephalin (as lock mass). DESI solvent was pumped using a micro Binary Solvent Manager (µBSM, Waters Corporation) at 3 µL/min flowrate. High performance sprayer for DESI utilized a capillary voltage of 0.65-0.75 kV with a nebulizing nitrogen gas. The ions were shuttled inside the mass spectrometer using a heated transfer line kept at 100 °C. Data was acquired with Ion Mobility and all data were analyzed using MassLynx^TM^ (version 4.2, Waters Corporation), DriftScope ^TM^ (version 2.9, Waters Corporation), and HDI ^TM^ (version 1.6, Waters Corporation) software.

### Immunohistochemistry staining

Tissues were first fixed overnight in 10% NBF (neutral buffered formalin) and processed for paraffin embedding. Paraffin sections were boiled in Trilogy reagent (Millipore-Sigma) for 15 min at 250 °F in a pressure cooker to facilitate deparaffinization, rehydrating, and antigen unmasking. Sections were washed in PBS and treated with 3% hydrogen peroxide for 15 min to block peroxidases. Sections were washed in PBST (PBS with 0.1% Tween-20) and blocked in 5% horse serum/1% BSA for 1 h. Primary antibodies diluted in PBST with 5% horse serum/1% BSA were then added to the sections for incubation overnight at 4°C. For immunohistochemistry (IHC), antibodies were detected with the ImmPRESS horse anti-rabbits kit and DAB (Vector Labs), then counterstained with hematoxylin (Newcomer Supply). The tissue slides were dehydrated in increasing amounts of ethanol and finally xylenes (Millipore-Sigma). The primary antibody used was: Cleaved Caspase-3 (Cell Signaling Technology, 9664) (1:250). Abundance of CC3^pos^ cells in DMS79 xenograft after combinatorial treatment of venetoclax and Cys depletion was quantified using ImageJ.

For immunohistochemistry tissues were fixed in 4% Formaldehyde solution (Merck) and subsequently embedded in paraffin. After sectioning tissue, it was deparaffinized and rehydrated according to standard procedure. For CC3 staining the following staining protocol was used. Endogenous peroxidase activity was quenched using Bloxall solution (Vector Laboratories) for 15 min, followed by three washes in tap water (3 × 5 min). Heat-induced antigen retrieval was performed in Vector Antigen Retrieval Buffer (1:100 dilution) using a pressure cooker for 45 min. After cooling, slides were washed in PBS for 5 min. Non-specific binding was blocked by incubating the sections for 60 min in blocking buffer (1% BSA, 0.05% Tween-20, 0.2% fish gelatine, and 0.003% sodium azide in PBS), supplemented with Avidin (Vector Avidin/Biotin Blocking Kit). Sections were then incubated overnight at 4°C with anti-cleaved caspase-3 rabbit monoclonal antibody (Cell Signaling Technology, #9661, 1:500 dilution) diluted in blocking buffer supplemented with biotin and PBS. The following day, slides were washed in PBS with 0.05% Tween-20 (PBS-T; 3 × 5 min), then incubated for 60 min at room temperature with biotinylated anti-rabbit secondary antibody (PerkinElmer, NEF813, 1:1000) in blocking buffer. Subsequently, slides were incubated with the VECTASTAIN Elite ABC-HRP complex (Vector Laboratories, prepared 1:60 in PBS) for 30 min, followed by DAB staining (Abcam, ab6423) according to the manufacturer’s instructions.

For 4-HNE staining the following protocol was used. Paraffin-embedded tissue sections were stained for 4-hydroxynonenal (4-HNE) using a mouse monoclonal anti-4-HNE antibody (clone HNEJ-2, JaICA). Sections were washed in 1× TBS-T (0.01% Tween-20 in TBS) and blocked for 20 min at room temperature using TBS-T supplemented with 10% normal goat serum. After a brief wash in TBS-T, slides were incubated overnight at 4°C in a humidified chamber with the primary antibody (diluted 1:100 in TBS-T with 5% goat serum). The next day, sections were washed (3 × 2 min, TBS-T) and endogenous peroxidase activity was quenched using 0.3% hydrogen peroxide in methanol for 20 min. After another TBS-T wash (3 × 2 min), sections were incubated with a biotinylated goat anti-mouse secondary antibody (Biotin-SP (long spacer) AffiniPure® Goat Anti-Mouse IgG (H+L) 115-065-166, 1:200 dilution in TBS-T with 5% goat serum) for 30 min at room temperature. An avidin–biotin–peroxidase complex (ABC) was prepared using the VECTASTAIN Elite ABC Kit (Vector Laboratories, PK-6100) in PBS and incubated on slides for 30 min at room temperature. Following washes in TBS-T (3 × 5 min), DAB staining (Abcam, ab6423) was performed according to the manufacturer’s instructions.

Abundance of CC3^pos^ cells and DAB intensity (4-HNE) on tumor sections was quantified using QuPath.

### Dichlorodihydrofluorescein diacetate (DCF-DA) reactive oxygen species (ROS) assay

Cells were plated at 5 × 10^5^ cells/well in 6-well plates with Cys replete or depleted RPMI medium containing 2% BGS. At each time point cells were washed with Hanks’ Balanced Salt Solution (HBSS) (Thermo Fisher Scientific, 14185052), suspended in 2 mL of 100 µM DCF-DA in HBSS, and incubated at 37 °C for 30 min. After wash in HBSS, 2 × 10^5^ cells in 100 µL were plated in 96-well plate. Fluorescence was measured at excitation and emission wavelengths of 485 nm and 520 nm using a plate reader.

### IncuCyte cell imaging

Cells were plated in 96-well plates (1x 104 stickers, 2 x 104 floaters) one day prior treatment. Cells were imaged for 72 h every 2 h using the 10× objective within the IncuCyte SX5 live cell imaging system (Sartorius) and 4 images per well were captured. The next day, cells were incubated with desired combinations of drugs. To detect cell death, 100 nM DRAQ7 (Thermo Fisher Scientific) was added to all wells while lipid ROS accumulation was assessed by addition of 1 µM STY-BODIPY. For cleaved Caspase 3 (CC3) cells were co-incubated with 5 μM CellEvent^TM^ Caspase3/7 Detection Reagent (Invitrogen). Each condition was tested in three technical replicates. Levels of lipid peroxidation were monitored by quantifying levels of co-oxidized STY-BODIPY (green, λex = 488 nm, λem = 495−540 nm) emitting in the green spectrum over reduced STY-BODIPY (red, λex = 561 nm, λem = 568−630 nm) emitting in the orange spectrum. Analysis of the number of DRAQ7-positive (dead) cells, CC3-positive cells, as well as reduced and oxidized STY-BODIPY was performed with IncuCyte Software v2021A and v2022B (Sartorius).

### Cell viability assay

For AlamarBlue assays (Millipore-Sigma, R7017), cells were plated at 1 × 10^5^ cells/well in 96-well plates with treatments. After 24 h, 48 h, and 72 h of culture, cell viability was assessed by measuring fluorescence at excitation and emission wavelengths of 530 nm and 590 nm, respectively, using a plate reader.

To determine cell viability using CellTiter-Glo assay, 20,000 cells were seeded into the wells of a 96-well plate. Additionally, control wells containing medium without cells to obtain a value for background luminescence were prepared. The test compounds were added to each well and incubated for the desired time. Then, the plate was equilibrated at room temperature for approximately 30 min, followed by a transfer of the cells to a white 96-well plate suitable for luminescent measurements. A volume of CellTiter-Glo reagent (Promega, G9242) equal to the volume of cell culture medium present in each well was added. Contents were mixed for 2 min on an orbital shaker to induce cell lysis, followed by incubation at room temperature for 10 min to stabilize luminescent signal. Luminescence signal was detected using a Tecan plate reader.

### Cysteinase kinetics assay

Cysteine kinetics assay was conducted as previously described^65^. Briefly, Eppendorf tubes containing 220 µL of cystine or blanks in 100 mM sodium phosphate buffer, 1 mM EDTA (pH 7.3), and 10 µM PLP were started by adding 20.3 µL of enzyme solution. After a set length of time, the reaction was quenched with 26.7 µL of 50% trichloroacetic acid. Then MBTH solution (2.2 mL: 1.6mL of 1M sodium acetate buffer pH 5.0 and 0.6 mL of 0.1% MBTH in same) was added and incubated at 50°C for 1 h. 100 µL of the solution was transferred to 96-well plate and absorbance at 320 nm was determined using a plate reader.

### Enzyme-linked immunosorbent assay (ELISA)

Samples for ELISA were either processed directly after harvest or stored at -80°C. ELISAs were performed using the BH2 ELISA kit (MyBioSource, MBS2568027) following the manufacturer’s protocol. Absorbance was measured at 450 nm with Multiskan Sky microplate spectrophotometer. The average absorption of 0 pg/mL standard was subtracted from absorption values of other standards and samples. Results were calculated by plotting a four-parameter logistic curve using GraphPad Prism.

### Gene expression analysis in SCLC cell lines and patient tissue

To determine the expression levels of genes relevant to the ferroptosis and BH2/BH4 synthesis we used public data as in^120^. To determine expression levels of *GNMT* and *SLC7A11* in SCLC tumors in comparison to adjacent lung tissue, we used public data as in ^70^. To determine expression levels *GCH1* and *ASCL1* in human SCLC patient tissue, we used public data from ^44^. UMAP projections from *RPM*-derived SCLC cell lines were generated using public data from ^15^. UMAP projections from human SCLC patient data were generated using public data from ^7^. Published Chip-Seq datasets of human SCLC cell lines^77-80^ for binding of ASCL1 and NEUROD1 were retrieved from ChiP-Atlas.org in the form of bigWig files, imported into the Integrative Genomics Viewer (IGV_2_17.4, Broad Institute) for visualization and aligned to hg38 in the promoter region of GCH1 on chromosome 14.

### Statistical analysis

Statistical significance was assayed with GraphPad Prism software (version 10). The specific tests used are indicated in the figure legends. No data were excluded from the experiments presented. Mice for *in vivo* studies were randomized where applicable and sample sizes determined by pilot experiments or previous experiments using similar models. Investigators were not blinded to allocation during experiments and outcome assessment.

### Data availability

The RNA-seq data for the RPR2 model are available at the Gene Expression Omnibus (GEO, accession number GSE171919). All other genomic and metabolic datasets generated in this study will be made available. All other data are available in the article and supplementary materials, or from the corresponding author upon reasonable request.

## ACKNOWLEDGEMENTS

We thank all the members of the Sage and von Karstedt labs for their help and support throughout this study. We thank Bandana Prasad from Agilent Technology Inc. for assistance with RNA-seq data analysis using Agilent GeneSpring software. We thank the Metabolomics Knowledge Center at Stanford Sarafan ChEM-H for providing 6545 Q-TOF LC-MS instrumentation and collaboration. We thank the Stanford University Mass Spectrometry for assistance with *in situ* metabolomics.

## AUTHOR CONTRIBUTIONS

J.W.K., C.M.B., S.v.K. and J.S. designed most of the experiments and interpreted the results; Y.D. and J.W.K, M.Y., D.P. and C.M.B. performed the metabolomic experiments together under the guidance of C.F.; J.W.K. and C.M.B. performed most of the experiments with help from S.B., A.E., T.R., L.B., M.B., T.N., Je.S., Y.T.S., Y.N., K.S., A.P.D., and A.C.C.; A.T.A. analyzed RNA-seq data; A.L. and Y.L. performed the metabolic tracing with J.W.K. under the guidance of J.Y.; Y.P.K. performed assays related to cysteinase activity with J.W.K. under the guidance of G.M.D.; C.M.B and L.B.L. analyzed cell death mechanisms in SCLC cells with J.W.K. under the guidance of S.v.K. and S.J.D.; H.J.O. and B.S. performed the analysis of metabolites on SCLC tumor sections; P.M., A.Q.V., and C.M.R. provided molecularly-characterized PDX models; J.W.K., C.M.B., S.v.K and J.S. wrote the manuscript and prepared the figures with contributions from all authors.

## FIGURES AND FIGURE LEGENDS

**Figure S1, related to Figure 1:**
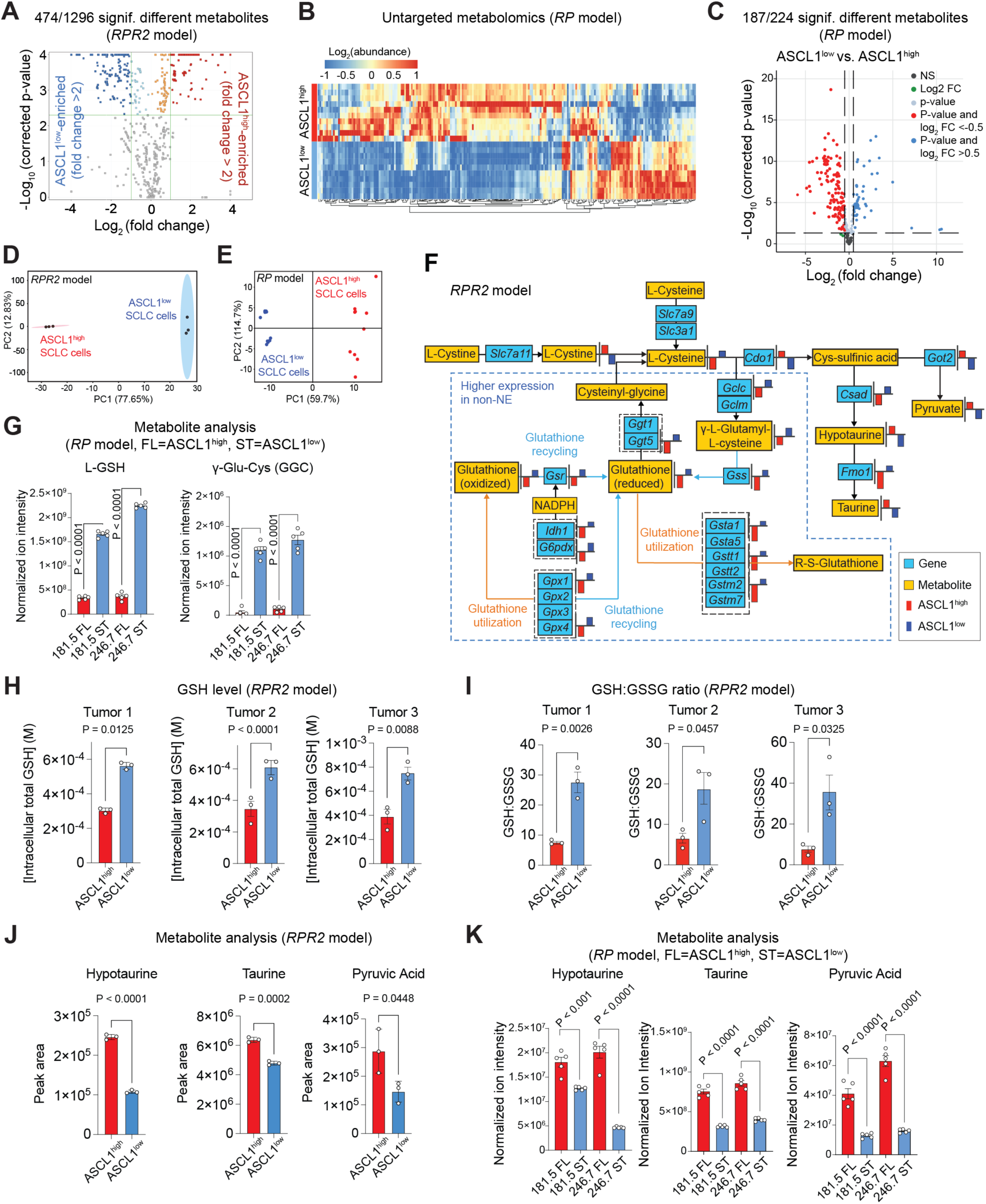
Metabolomic analyses reveal differentially regulated metabolites between ASCL1^high^ and ASCL1^low^ mouse SCLC cells. **A.** Volcano plot showing differentially regulated metabolites in ASCL1^high^ and ASCL1^low^ mouse SCLC cells from *RPR2* mutant tumors (as in Figure 1A,B) (n=3). -Log10(p-value) is capped at 4 for display. **B.** Hierarchical clustering analysis of untargeted metabolomics in the *RP* model. **C.** Volcano plot showing differentially regulated metabolites in ASCL1^high^ and ASCL1^low^ mouse SCLC cells from *RP* mutant tumors (as in Figure 1A) (n=3). **D.** Principal component analysis (PCA) of metabolites in ASCL1^high^ and ASCL1^low^ mouse SCLC cells in the *RPR2* model. **E.** Principal component analysis (PCA) of metabolites in ASCL1^high^ and ASCL1^low^ mouse SCLC cells in the *RP* model. **F.** Analysis of the Cys-glutathione pathway in ASCL1^high^ and ASCL1^low^ mouse SCLC cells in the *RPR2* model. Differentially regulated genes and metabolites are in light blue and yellow, respectively. Bar graphs represent Log2 of normalized (to baseline using mean) values of FPKM for genes and of peak area for metabolites. **G.** Abundance of GSH and y-Glu-Cys detected by LC/MS in ASCL1^high^ and ASCL1^low^ mouse SCLC cells in the *RP* model (n=5). **H.** Intracellular total glutathione (GSH) concentration in 5x10^6^ ASCL1^high^ and ASCL1^low^ mouse SCLC cells (in 100 µL) quantified using a colorimetric assay (n=3) (*RPR2* model). **I.** Intracellular ratio of GSH and glutathione disulfide (GSSG) concentration in ASCL1^high^ and ASCL1^low^ mouse SCLC cells quantified using a colorimetric assay (n=3) (*RPR2* model). **J.** Abundance of hypotaurine, taurine, and pyruvic acid detected by LC/MS in ASCL1^high^ and ASCL1^low^ mouse SCLC cells (n=3) (*RPR2* model). **K.** Abundance of hypotaurine, taurine, and pyruvic acid detected by LC/MS in ASCL1^high^ and ASCL1^low^ mouse SCLC cells (n=5) (*RP* model). P-values were calculated by Student’s t-test. Error bars indicate mean ± SEM.

**Figure S2, related to Figure 1:**
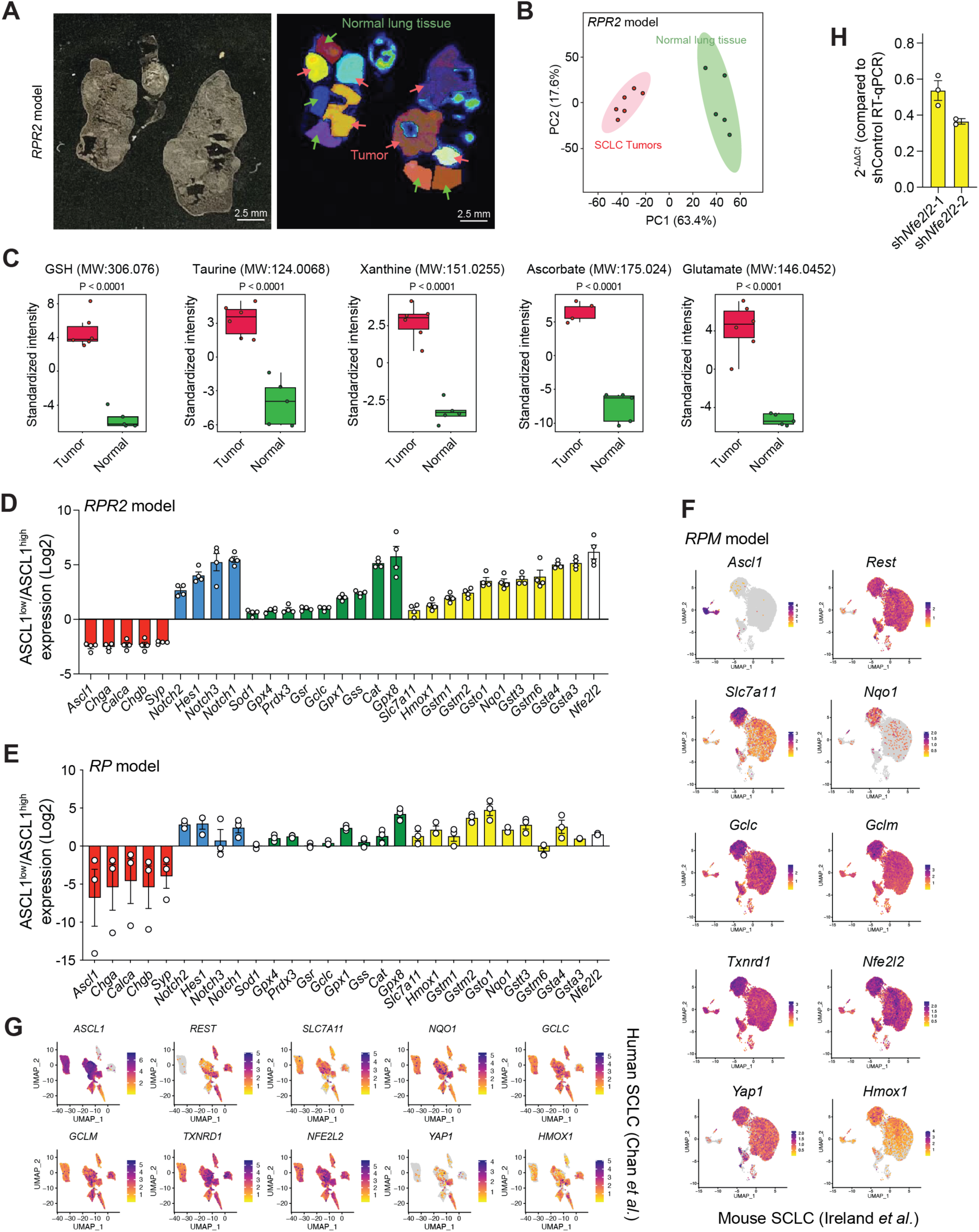
Metabolomic and transcriptomic analysis of NRF2 and glutathione in ASCL1^high^ and ASCL1^lowh^ mouse SCLC cells. **A.** Representative images of a cryosection slide (left) and spatially clustered regions (differently colored) from mass spectrometry imaging (right, colored) from the lungs of *RPR2* mutant mice. Scale bar, 2.5 mm. **B.** Principal component analysis (PCA) of spatially clustered regions in (A) (red: tumors, green: normal tissue). **C.** Standardized intensity of selected metabolites detected by mass spectrometry imaging in SCLC tumors (red) and normal lung tissue (green) (n=5-6 regions, from n=1 *RPR2* mutant mouse). **D.** Expression levels (RNA-seq) of neuroendocrine (red) and non-neuroendocrine (blue) markers and antioxidant-response-element containing genes that are regulated by NRF2 (green: antioxidant genes; yellow: detoxifying genes; white: *Nfe2l2*) in ASCL1^low^ relative to ASCL1^high^ mouse SCLC cells (n=4) (*RPR2* model). **E.** Expression levels (RNA-seq) of neuroendocrine (red) and non-neuroendocrine(blue) markers and antioxidant-response-element containing genes that are regulated by NRF2 (green: antioxidant genes; yellow: detoxifying genes; white: *Nfe2l2*) in ASCL1^low^ relative to ASCL1^high^ mouse SCLC cells (n=4) (*RP* model). **F.** UMAP projection of a scRNA-seq dataset of mouse SCLC cells undergoing a neuroendocrine to non-neuroendocrine transdifferentiation showing expression of NRF2 target genes (from ^15^). **G.** UMAP projection of a scRNA-seq dataset of human SCLC cells showing expression of NRF2 target genes (from ^7^). **H.** RT-qPCR validation of *Nfe2l2* knockdown in ASCL1^low^ mouse SCLC cells compared to control cells (n=3). Error bars indicate mean ± SEM.

**Figure S3, related to Figure 2:**
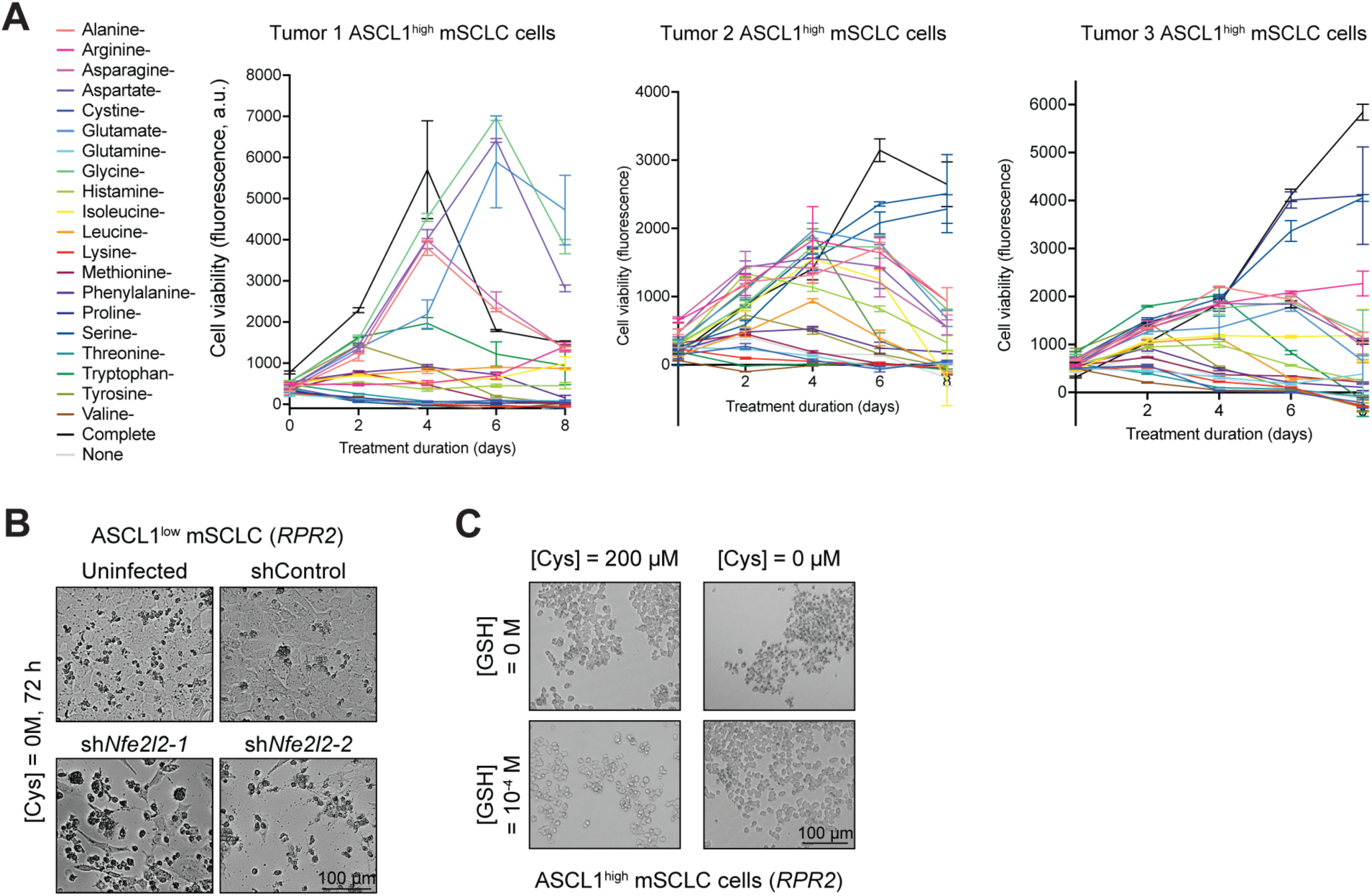
SCLC cells can be targeted by exogenous Cys deprivation in the culture medium. **A.** Proliferation of ASCL1^high^ mouse SCLC cell lines (mSCLC) in amino acid-depleted media over an 8-day time course (n=3). Cell viability on each day was measured as fluorescence values with arbitrary unit (a.u.) from an AlamarBlue assay. **B.** Representative images of ASCL1^low^ mSCLC cells with stable knock-down of *Nfe2l2* (sh*Nfe2l2*) (coding for NRF2) or GFP (as a control, shControl) and cultured in Cys-depleted medium for 72 h (see Figure 2C for quantification of cell death). Scale bar, 100 μm. **C.** Representative images of ASCL1^high^ mouse SCLC cells supplemented with GSH [100 µM] and cultured in complete medium (with 200 µM of Cys) or Cys-depleted medium for 72 h (see Figure 2D for quantification of cell death). Scale bar, 100 μm.

**Figure S4, related to Figure 2:**
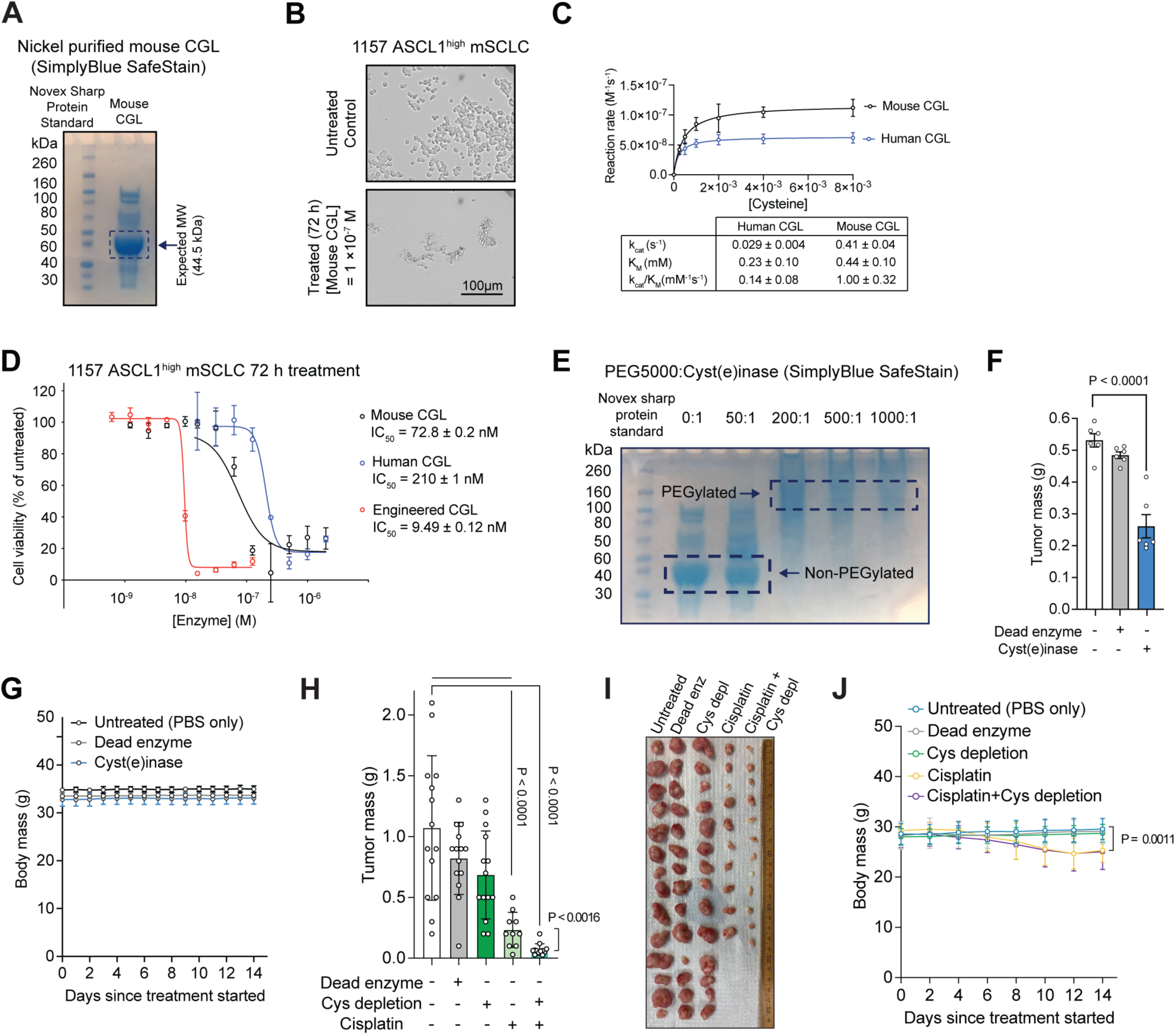
SCLC cells can be targeted by Cys deprivation in mice. **A.** Sodium dodecyl sulfate-polyacrylamide gel electrophoresis (SDS-PAGE) of His-tagged mouse cystathionine γ-lyase (CGL) purified using Nickel-Nitriloacetic acid (Ni-NTA). **B.** ASCL1^high^ mouse SCLC cells (mSCLC) were treated with mouse CGL [100 nM] and photographed 72 h after the treatment began. Scale bar, 100 µm. **C.** Michaelis-Menten curves and parameters of human and mouse CGL. Results are from experiments performed in triplicate. **D.** Dose-response curves and IC50 values of ASCL1^high^ mSCLC cells treated with mouse CGL, human CGL, and engineered CGL measured by cell viability (AlamarBlue assay) 72 h after the treatments began (n=3). **E.** SDS-PAGE of engineered CGL conjugated with different molar ratios with PEG5000. **F.** Final time point tumor mass measurements as in Figure 2H. **G.** Body mass of mice as in Figure 2H. **H.** Final time point tumor mass measurements as in Figure 2K. **I.** Image of collected tumors at final time point as in Figure 2K. **J.** Body mass of mice as in Figure 2K. P-values were calculated by one-way ANOVA (P<0.0001) followed by the unpaired Student’s t-test in (F), (H), and (J). Error bars for (C) and (D) indicate mean ± SD. Error bars for all other figures indicate mean ± SEM.

**Figure S5, related to Figure 3:**
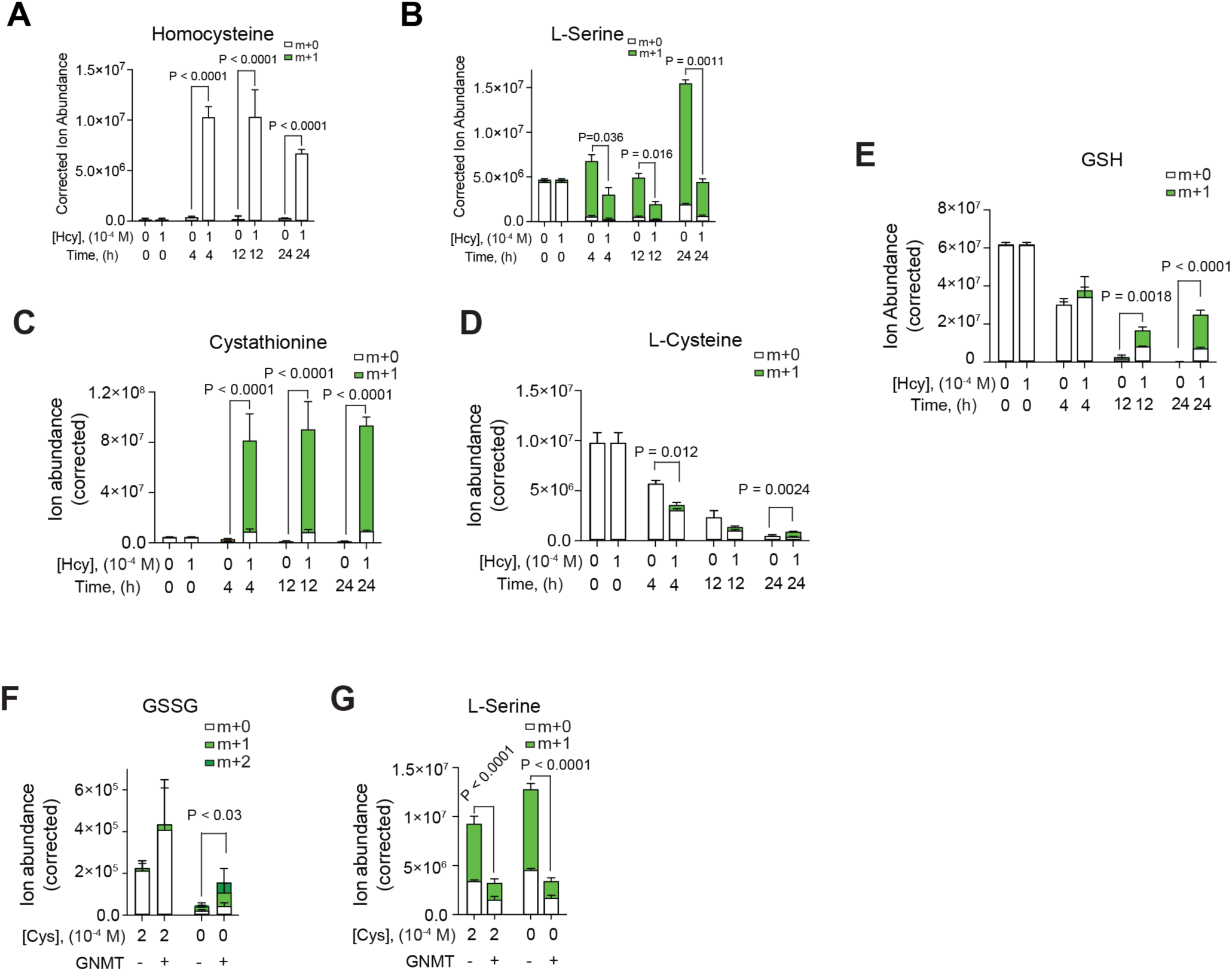
SCLC cells can be targeted by Cys deprivation in mice. **A-E.** Labelled and unlabeled homocysteine (A), L-serine (B), cystathionine (C), L-cysteine (D, and GSH (E) in NCI-H82 SCLC cells (NEUROD1^high^) growing in Cys replete medium containing ^13^C-labeled L-serine supplemented with homocysteine [100 µM] for 24 h (n=2). **F,G.** Labeled and unlabeled GSSG (F), and L-serine (G) in NCI-H82 SCLC cells (NEUROD1^high^) ectopically expressing GNMT in Cys replete and depleted medium (n=4). P-values were calculated by the unpaired Student’s t-test. Error bars indicate mean ± SEM.

**Figure S6, related to Figure 4:**
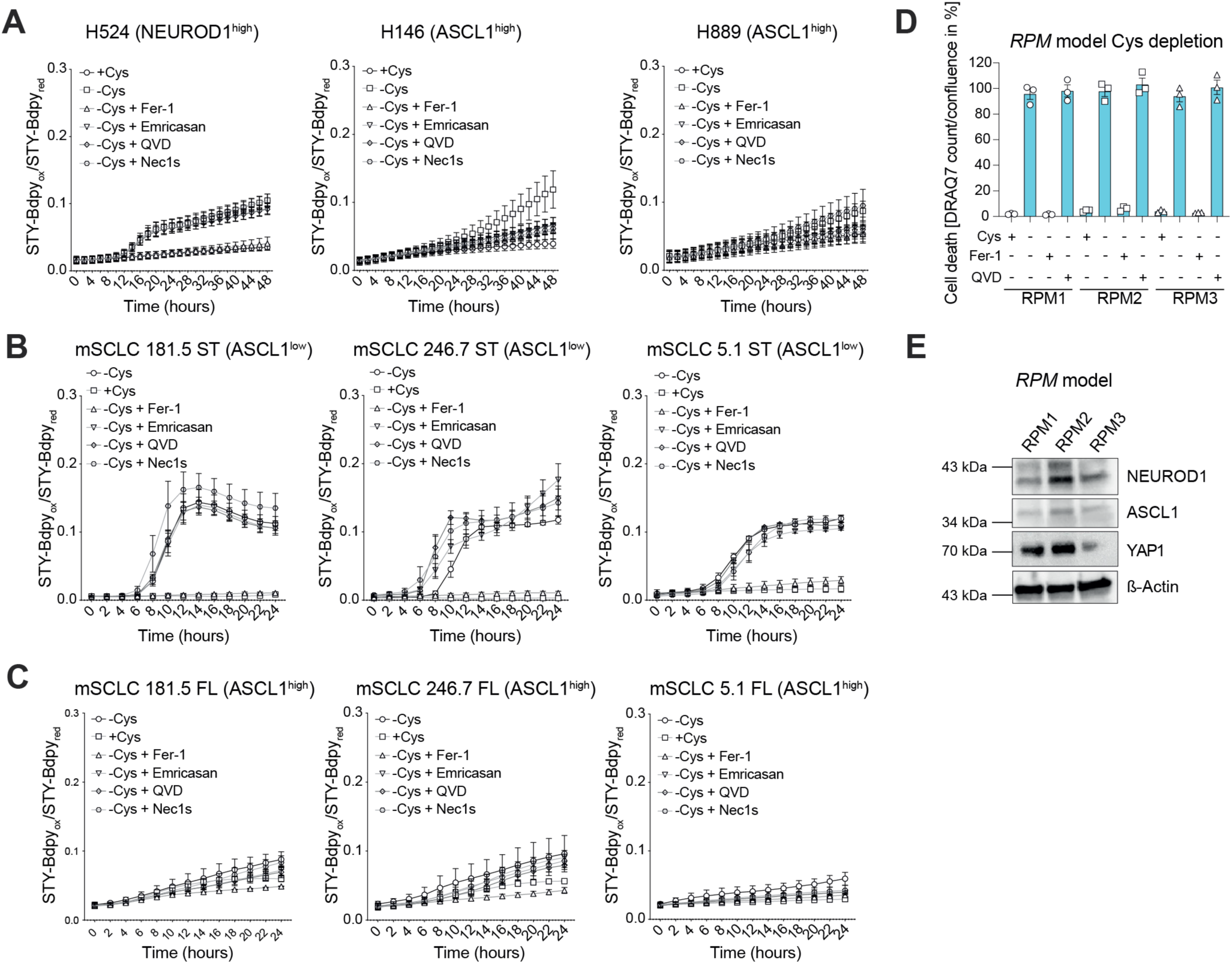
Cys deprivation induces ferroptosis in ASCL1^low^ SCLC cells. **A.** Human SCLC cell lines as indicated were cultured in Cys-depleted medium, supplemented with Cys [200 µM], Fer-1 [1 µM], Emricasan [2.5 µM], QVD [10 µM] and Nec1s [10 µM] and stained for lipid ROS accumulation using STY-BODIPY [1 μM] for 48 h (n=3). **B,C.** *RP*-derived mouse ASCL1^low^ (ST) (B) and ASCL1^high^ (FL) (C) SCLC cell lines (mSCLC) were cultured in Cys-depleted medium, supplemented with Cys [200 µM], Fer-1 [1 µM], Emricasan [2.5 µM], QVD [10 µM] and Nec1s [10 µM] and stained for lipid ROS accumulation using STY-BODIPY [1 μM] for 24 h (n=3). **D.** Cell death (DRAQ7 count/confluence in %) of *RPM*-derived SCLC cell lines (RPM1, -2, -3) cultured in Cys-depleted medium, supplemented with Cys [200 µM], Fer-1 [1 µM] and QVD [10 µM] for 24 h. **E.** Representative Immunoassays (n=3) of ASCL1^low^ *RPM*-derived SCLC cell lines as in (e). ß-Actin was used as a loading control.

**Figure S7, related to Figure 5:**
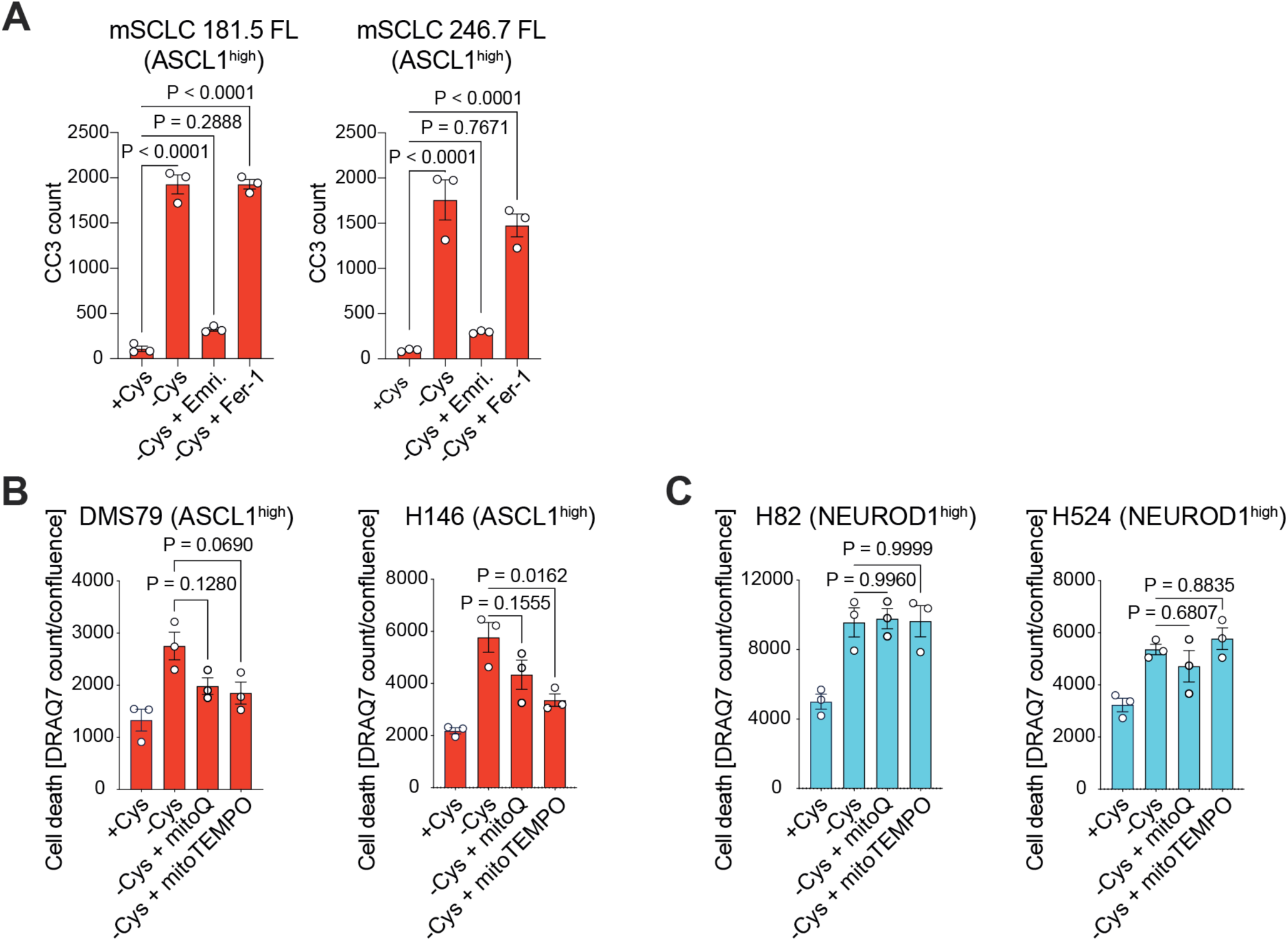
ASCL1^high^ SCLC cells undergo apoptosis in response to Cys deprivation. **A.** *RP*-derived ASCL1high mSCLC cells (181.5 FL, 246.7 FL) were cultured in Cys-depleted medium supplemented with Cys [200 µM], Emricasan (Emri.) [2.5 µM] and Fer-1 [1 µM] for 48 h. Caspase activity as a sign of apoptotic cell death (CC3 count) was monitored using a fluorogenic real-time reporter measured using the IncuCyte SX5 bioimaging platform. **B,C.** Cell death (DRAQ7 count/confluence) of human SCLC cell lines (ASCL1^hig^h in B, NEUROD1^high^ in C) cultured in Cys-depleted medium supplemented with Cys [200 µM], mitoquinone (MitoQ) [50 nM], and MitoTempo [2.5 µM] as indicated for 72 h. For (A,B,C) one-way ANOVA was performed, and Tukey was used as post hoc test. Error bars indicate mean ± SEM.

**Figure S8, related to Figure 6:**
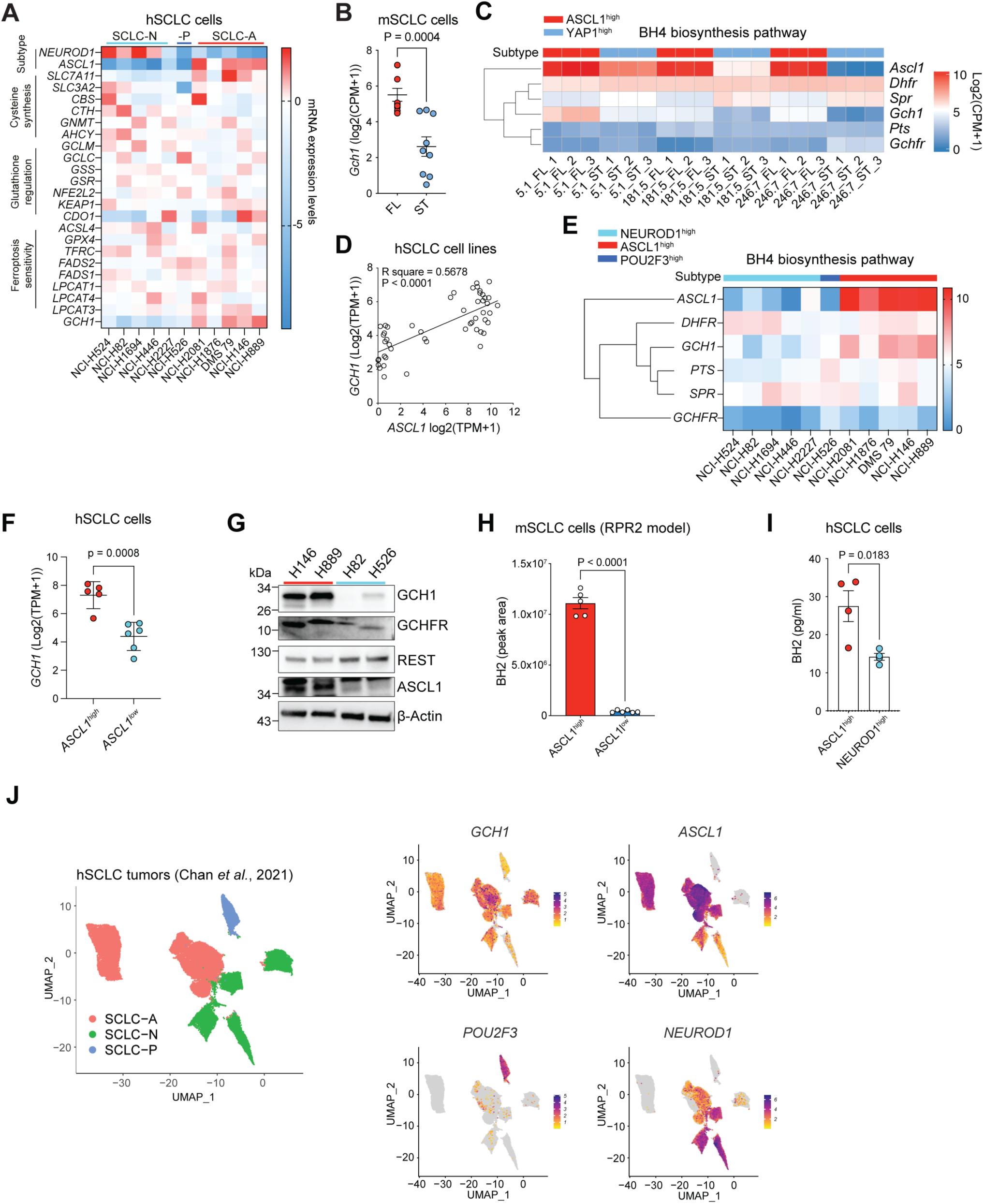
The BH4 biosynthetic pathway is upregulated in ASCL1^high^ SCLC cells. **A.** Heatmap of gene expression levels associated with *de novo* cysteine synthesis, glutathione regulation, and ferroptosis sensitivity in selected human SCLC (hSCLC) cell lines (RNA levels data from the EMBL-EBI Expression Atlas). **B.** RNA-seq data from three different *RP*-derived mouse ASCL1^high^ (FL) and ASCL1^low^ (ST) SCLC cell lines (mSCLC) (181.5, 246.7, and 5.1) plotted as log2(CPM+1) for *Gch1* expression (technical triplicates for each cell line). **C.** Heatmap of gene expression levels associated with the BH2/BH4 biosynthesis pathway in the cell lines as in (B). **D.** Scatter plots of RNA-seq expression data from hSCLC cell lines (Expression Atlas) for *GCH1* and *ASCL1*. **E.** Heatmap of gene expression levels associated with BH4 biosynthesis, *ASCL1*, *NEUROD1*, and *POU2F3* in selected hSCLC cell lines (RNA levels data from the EMBL-EBI Expression Atlas). **F.** Data from (e) for *GCH1* in ASCL1^high^ and ASCL1^low^ hSCLC cells. **G.** Representative immunoassay of hSCLC cell lines for GCH1, GCHFR, REST and ASCL1. β-Actin was used as a loading control. **H.** Abundance of BH2 detected by LC/MS in *RPR2*-derived mouse ASCL1^high^ and ASCL1^low^ SCLC cells (n=5-6). **I.** Abundance of BH2 quantified using ELISA in four ASCL1^high^ cell lines and three NEUROD1^high^ human cell lines. **J.** UMAP projection of a published dataset from human SCLC patient data showing classification into the different SCLC subtypes by marker expression (data from ^7^) (left). UMAP projection of the same dataset showing expression of *GCH1, ASCL1, POU2F3, NEUROD1* genes. For (B,F,H,I) Student’s t-test was performed. Error bars indicate mean ± SEM.

**Figure S9, related to Figure 6:**
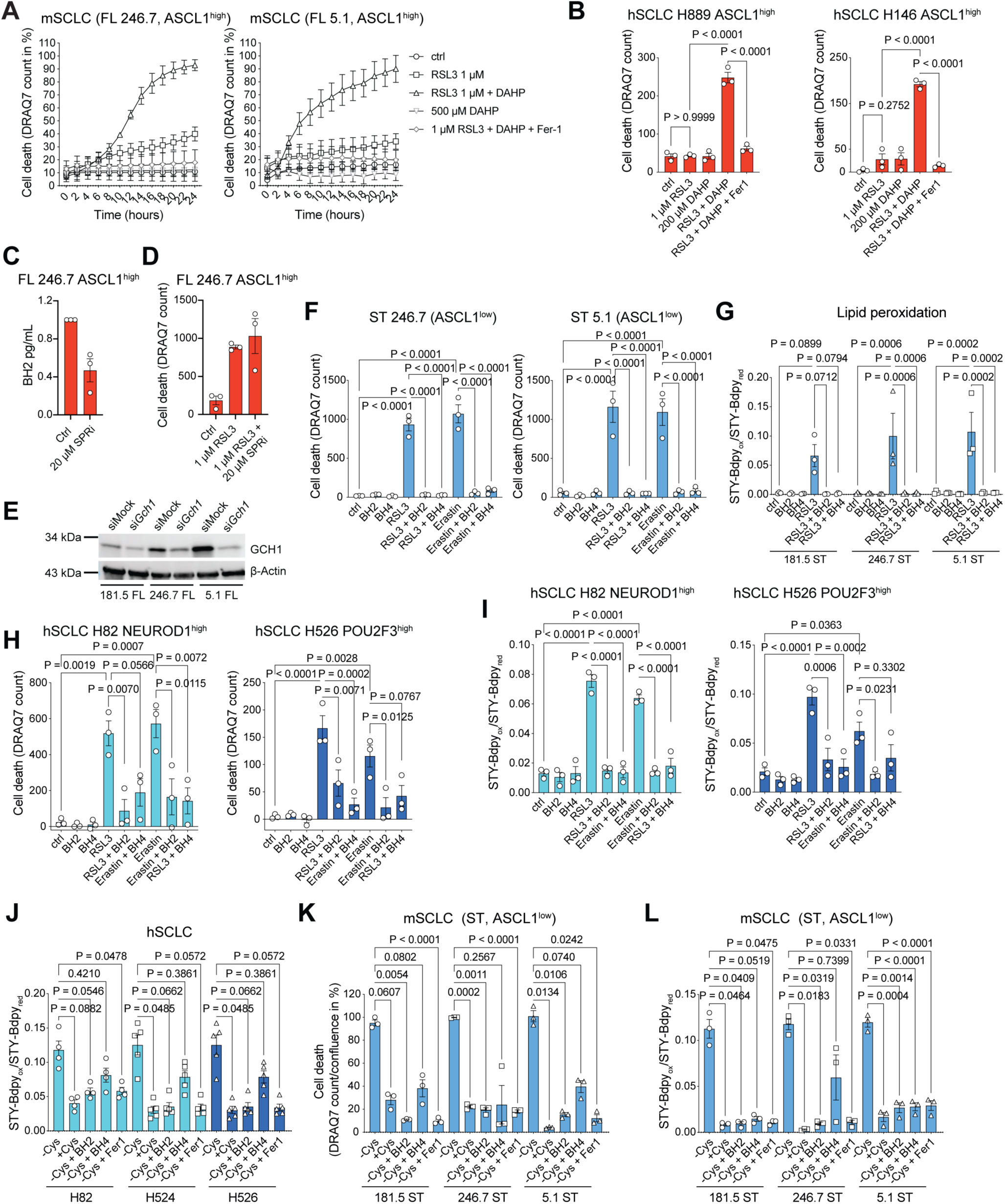
Upregulation of the BH4 biosynthetic pathway causes ferroptosis resistance in ASCL1^high^ SCLC cells. **A.** *RP*-derived mouse SCLC cells (246.7 FL, 5.1 FL) were cultured in full medium, treated with ferroptosis inducer RSL3 [1 µM], DAHP [500 µM], RSL3+DAHP and RSL3+DAHP+Fer-1 [1 µM] for 24 h. Cell death (DRAQ7 count in %) was monitored by DRAQ7 [100 mM] incorporation and measured using the IncuCyte SX5 bioimaging platform (n=3). **B.** hSCLC cell lines (H889, H146) were treated with RSL3 [1 µM], DAHP [200 µM], RSL3+DAHP and RSL3+DAHP+Fer-1 [1 µM] for 24 h. Cell death (DRAQ7 count) was monitored by DRAQ7 [100 mM] incorporation and measured using the IncuCyte SX5 bioimaging platform (n=3). **C.** *RP*-derived mouse SCLC cells (246.7 FL) were treated with Spri3 (SPR-inhibitor) [20 µM] for 24 h and BH2-levels were determined using ELISA (n=3). **D.** *RP*-derived mouse SCLC cells (246.7 FL) were treated with RSL3 [1 µM], Spri3 [20 µM] or RSL3+Spri3 for 24 h. Cell death (DRAQ7 count) was monitored by DRAQ7 [100 mM] incorporation and measured using the IncuCyte SX5 bioimaging platform. (n=3) **E.** Representative immunoassay (n=3) for GCH1 in ASCL1^high^ *RP-*derived mouse SCLC cells (181.5 FL, 246.7 FL, 5.1 FL) subjected to GCH1-knockdown (si*Gch1*) or control (siMock) for 5 days. ß-Actin was used as a loading control. **F.** *RP*-derived mouse SCLC cells (246.7 and 5.1 ST) were treated with RSL3 [1 µM], BH2 [50 µM], BH4 [100 µM], Erastin [10 µM] and the combination of each for 24 h. Cell death (DRAQ7 count) was monitored by DRAQ7 [100 mM] incorporation and measured using the IncuCyte SX5 bioimaging platform (n=3). **G.** *RP*-derived mouse SCLC cells (181.5, 246.7 and 5.1 ST) were treated as in (F) and stained for lipid ROS accumulation using STY-BODIPY [1μM] for 24 h (n=3). **H.** Cell death (DRAQ7 count) of hSCLC cell lines (H82 and H526) treated with BH2 [50 µM], BH4 [100 µM], RSL3 [1 µM], Erastin [10 µM] and the combination of each of them (n=3). **I.** hSCLC cell lines (H82 and H526) were treated as in (G). and stained for lipid ROS accumulation using STY-BODIPY [1μM] for 24 h (n=3). **J.** hSCLC cells (H82, H524, H526) were cultured in Cys-depleted medium, supplemented with Cys [200 µM], BH2 [50 µM], BH4 [100 µM] and Fer-1 [1 µM] and stained for lipid peroxidation using STY-BODIPY [1 μM] for 24 h (n=4-5). **K.** *RP*-derived mouse SCLC cell lines (181.5, 246.7, 5.1 ST) were cultured in Cys-depleted medium, supplemented with Cys [200 µM], BH2 [50 µM], BH4 [100 µM] and Fer-1 [1 µM]. Cell death (DRAQ7 count) was monitored by DRAQ7 [100 mM] incorporation and measured using the IncuCyte SX5 bioimaging platform (n=3). **L.** *RP*-derived mouse SCLC cell lines (181.5, 246.7, 5.1 ST) were treated as in (J). and stained for lipid peroxidation using STY-BODIPY [1 μM] for 24 h (n=3). For (B,F,G,H,I,J,K,L) one-way ANOVA was performed, Tukey was used as post hoc test. Error bars indicate mean ± SEM.

**Figure S10, related to Figure 6:**
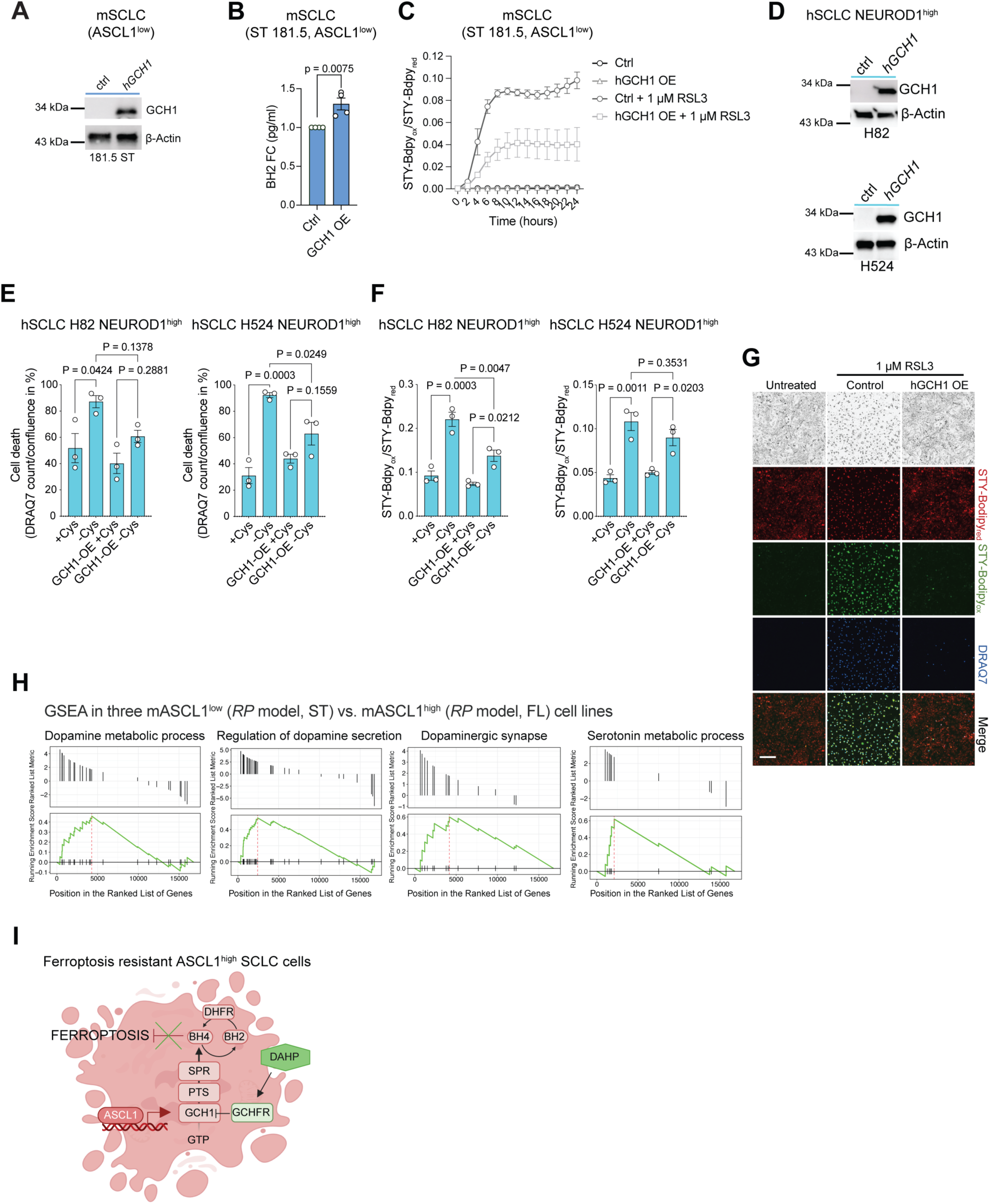
GCH1 expression is sufficient to render ASCL1^low^ SCLC cells ferroptosis resistant. **A.** Representative immunoassay (n=) for GCH1 in control *RP*-derived mouse SCLC cells (181.5 ST) and with hGCH1 overexpression for 24 h. ß-Actin was used as a loading control. **B.** Abundance of BH2 was measured by ELISA in *RP*-derived mouse SCLC cells (181.5 ST) subjected to hGCH1 overexpression for 24 h (n=4). **C.** *RP*-derived mouse SCLC cells (181.5 ST) were subjected to hGCH1 overexpression for 24 h and subsequently treated with RSL3 [1 µM] and stained for lipid peroxidation using STY-BODIPY [1 μM] for 24 h (n=3). **D.** Representative immunoassay (n=3) for GCH1 in control hSCLC cells (H82, H524) and with hGCH1 overexpression for 24 h. ß-Actin was used as a loading control. **E.** hSCLC cells (H82, H524) were subjected to hGCH1 overexpression for 24 h and subsequently cultured in Cys-depleted medium for 72 h. Cell death (DRAQ7 count/confluence in %) was monitored by DRAQ7 [100 nM] incorporation and measured using the IncuCyte SX5 bioimaging platform (n=3). **F.** hSCLC cells (H82, H524) were treated as in (E). and stained for lipid peroxidation using STY-BODIPY [1 μM] for 72 h (n=3). **G.** Representative images of data as shown in Figure 6K. Scale bar = 200 µm. **H.** GSEA analysis was performed on three different *RP*-derived mouse SCLC cells (181.5 246.7 5.1) comparing ST vs. FL. **I.** Scheme of GCH1-mediated ferroptosis resistance in ASCL1^high^ SCLC. For (B) Student’s t-test was performed. For (E) one-way ANOVA was performed, and Tukey was used as post hoc test. Error bars indicate mean ± SEM.

**Figure S11, related to Figure 7:**
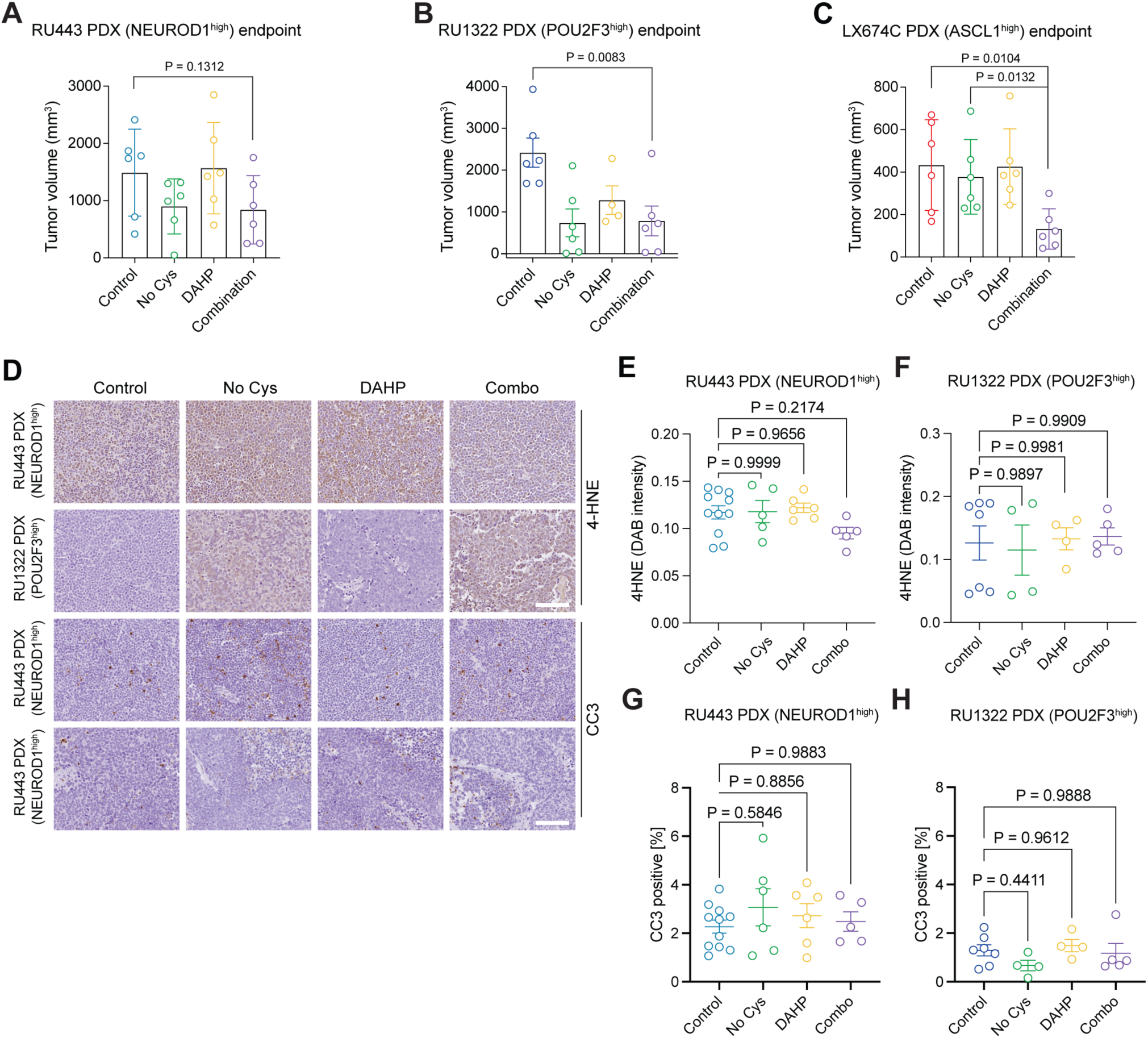
Cys depletion increases treatment susceptibility of SCLC patient-derived xenografts. **A.** Final volume measurements of the individual tumors in Figure 7A. **B.** Final volume measurements of the individual tumors in Figure 7B. **C.** Final volume measurements of the individual tumors in Figure 7C. **D.** Representative images of sections from RU1322 and RU443 tumors immunostained with cleaved caspase 3 (CC3) and 4-HNE antibodies. Scale bars, 100 µm. **E.** Quantifications of 4-HNE in sections from RU433 tumors (n(control)=11, n(No Cys)=6, n(DAHP)=6, n(combo)=5). **F.** Quantifications of 4-HNE in sections from RU1322 tumors (n(control)=7, n(No Cys)=4, n(DAHP)=4, n(combo)=5). **G.** Quantifications of CC3 in sections from RU433 tumors (n(control)=11, n(No Cys)=5, n(DAHP)=6, n(combo)=5). **H.** Quantifications of CC3 in sections from RU1322 tumors (n(control)=7, n(No Cys)=4, n(DAHP)=4, n(combo)=5). For (A, B, C) one-way ANOVA was performed followed by the unpaired Student’s t-test for post hoc analysis. For (E,F,G,H) one-way ANOVA was performed, and Tukey was used as post hoc test. Error bars indicate mean ± SEM.

**Figure S12, related to Figure 7:**
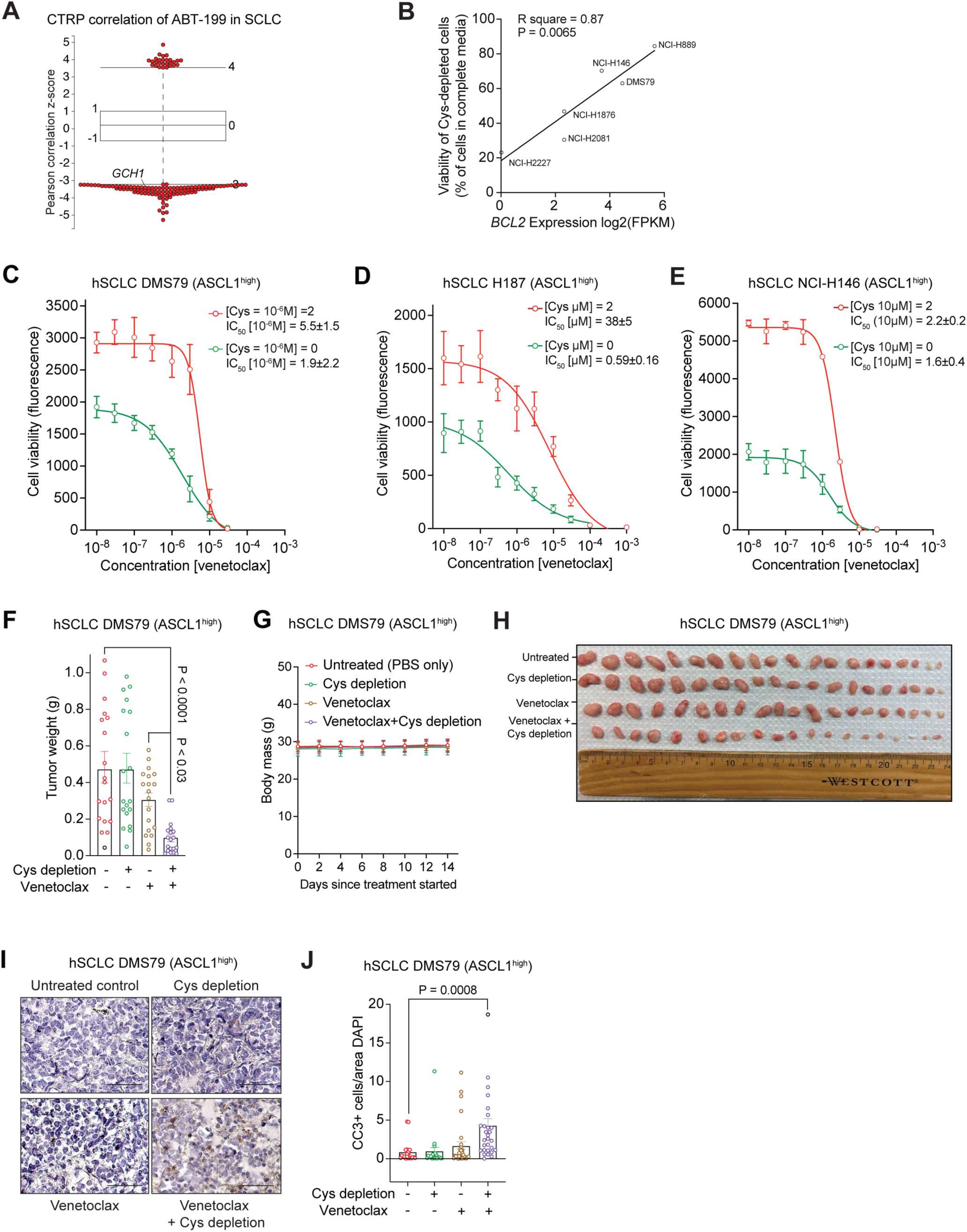
Cys depletion achieves superior tumor killing in combination with venetoclax. **A.** ABT-199 (venetoclax) sensitivity inversely correlates with expression of GCH1 in human SCLC cell lines. Plotted values are z-scored Pearson’s correlation coefficients; line, median; box, 25th to 75th percentile; whiskers, 2.5th and 97.5th percentile expansion Data from the CTRP database. **B.** Scatter plot of ferroptosis-resistant human SCLC cell lines displaying the correlation between viability under Cys deprivation and BCL-2 expression. **C-E.** Dose-response curves and IC50 values for NCI-H187 (c) and NCI-H146 (d) human ASCL1^high^ cells in Cys-replete or Cys-depleted medium treated with the BCL-2 inhibitor venetoclax, measured by cell viability (AlamarBlue assay) 72 h after the treatment began (n=3). **F.** Final time point tumor weight measurements as in Figure 7H. **G.** Body weights of mice as in Figure 7H. **H.** Image of collected tumors at final time point as in Figure 7H. **I.** Representative images of immunostained with a CC3 antibody (brown signal). Hematoxylin (blue) was used to counterstain Scale bars, 50 µm. **J.** Quantification of (I) (n=30). For (B) the Student’s t-test was used. For (F,J) one-way ANOVA was performed followed by the unpaired Student’s t-test for post hoc analysis.

## Supplementary Tables

**Table S1, related to Figure 1:** Pathways with at least 3 differentially regulated metabolites in mouse SCLC cells in the *RPR2* model. Note that individual metabolites in these pathways were not validated with standards for this analysis.

| Pathway | Matched Entitles | Pathway Entitles of Experiment Type |
| --- | --- | --- |
| Purine metabolism | 12 | 68 |
| tRNA charging pathway | 9 | 24 |
| Purine nucleotides degradation II (aerobic) | 8 | 19 |
| Oxidative stress and redox pathway | 8 | 24 |
| One carbon metabolism and related pathways | 8 | 41 |
| Purine and pyrimidine metabolism | 7 | 32 |
| Sphingolipid metabolism | 6 | 66 |
| Superpathway of citrulline metabolism | 6 | 30 |
| Gamma-glutamylcycle | 5 | 13 |
| Purine ribonucleosides degradation to ribose-1-phosphate | 5 | 13 |
| Sphingolipid metabolism | 5 | 26 |
| Citrulline biosynthesis | 4 | 22 |
| Glutathione and one carbon metabolism | 4 | 20 |
| Asparagine biosynthesis I | 4 | 9 |
| Arginine biosynthesis IV | 4 | 22 |
| Guanosine nucleotides degradation III | 4 | 12 |
| Eicosanoid metabolism via lipo oxygenases | 4 | 40 |
| Creatine biosynthesis | 4 | 14 |
| Glutathione biosynthesis | 4 | 9 |
| Kennedy pathway | 4 | 19 |
| Glutathione metabolism | 4 | 18 |
| Bile acid biosynthesis, neutral pathways | 4 | 48 |
| Citrulline-nitric oxide cycle | 3 | 13 |
| TCA cycle | 3 | 23 |
| Adenosine nucleotides degradation II | 3 | 13 |
| Rate biosynthesis/inosine 5'-phosphate degradation | 3 | 11 |
| Nicotine degradation III | 3 | 26 |
| Guanine and guanosine salvage II | 3 | 7 |
| Porphyrin and chlorophyll metabolism | 3 | 124 |
| Glycerolipids and glycerophospholipids | 3 | 29 |
| NAD biosynthesis II (from tryptophan) | 3 | 24 |
| Pyrimidine ribonucleotides de novo biosynthesis | 3 | 24 |
| Phosphatidylethanolamine biosynthesis II | 3 | 10 |
| L-carnitine biosynthesis | 3 | 14 |
| Nicotine degradation II | 3 | 27 |
| L-ascorbate biosynthesis VI | 3 | 19 |
| Ureide biosynthesis | 3 | 17 |
| Ceramide biosynthesis | 3 | 11 |
| Urea cycle | 3 | 17 |
| Glycine degradation (creatine biosynthesis) | 3 | 8 |

**Table S2, related to Figure 1:**
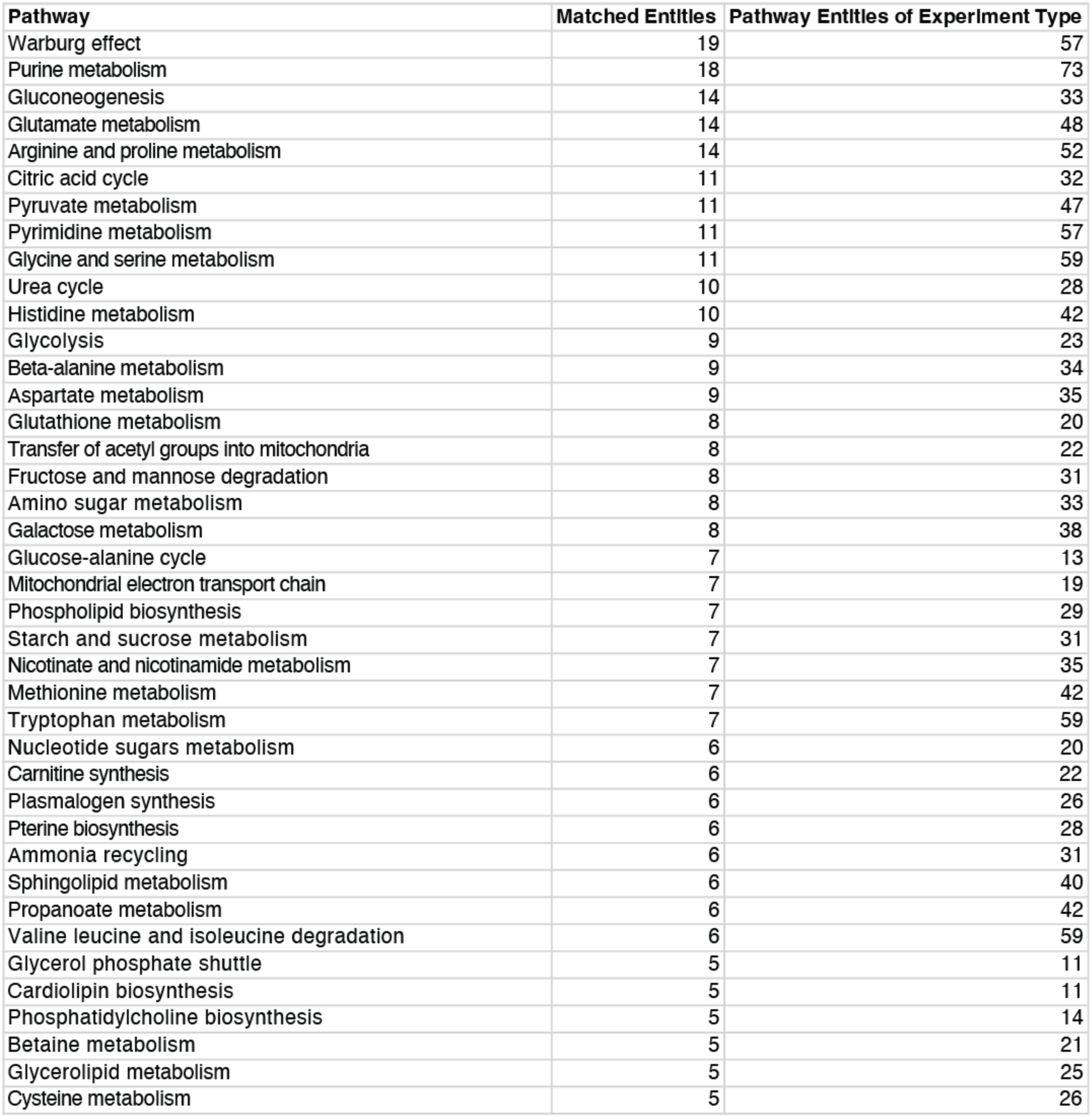
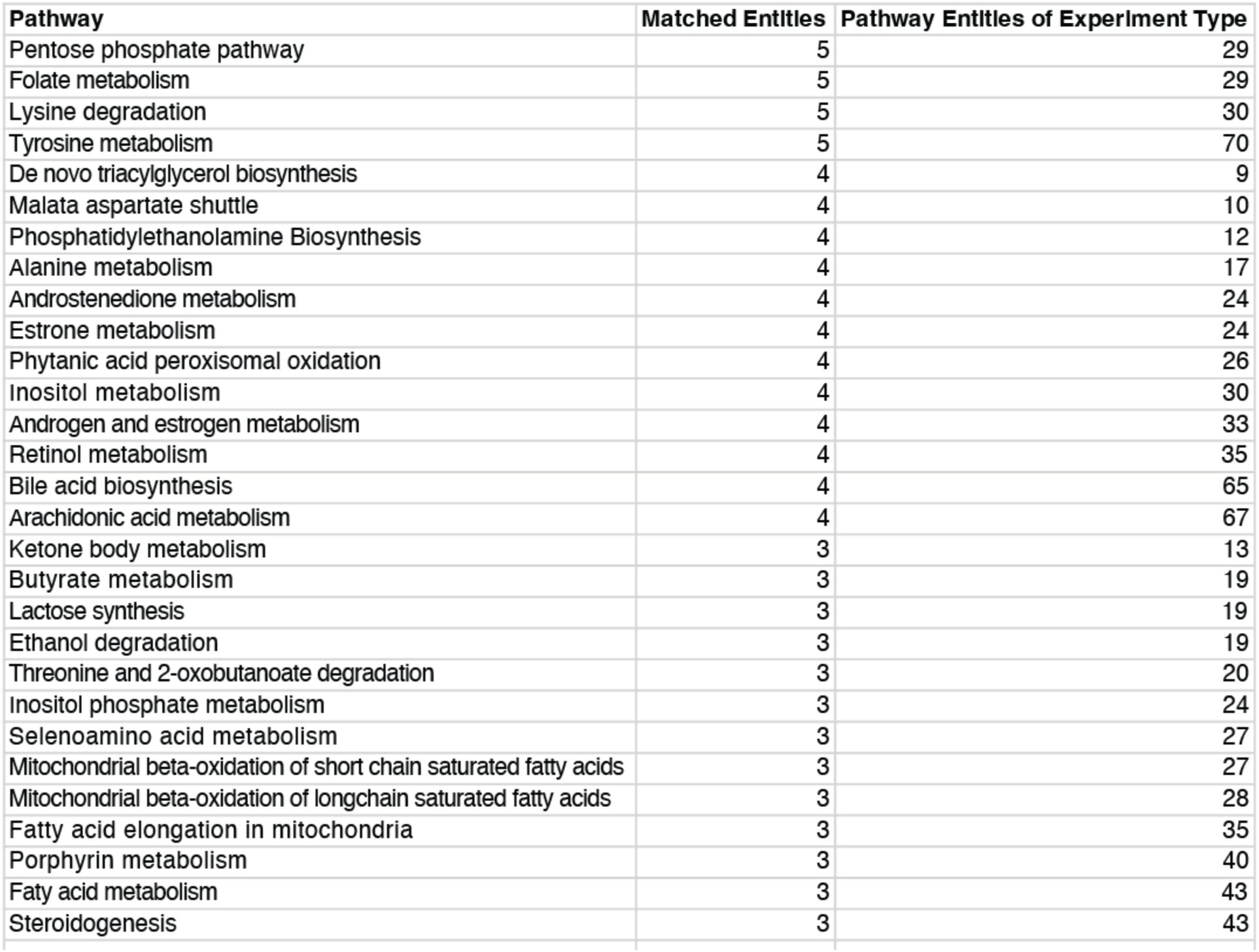
Pathways with at least 3 differentially regulated metabolites in mouse SCLC cells in the *RP* model. Note that individual metabolites in these pathways were not validated with standards for this analysis.

| Pathway | Matched Entitles | Pathway Entitles of Experlment Type |
| --- | --- | --- |
| Warburg effect | 19 | 57 |
| Purine metabolism | 18 | 73 |
| Gluconeogenesis | 14 | 33 |
| Glutamate metabolism | 14 | 48 |
| Arginine and proline metabolism | 14 | 52 |
| Citric acid cycle | 11 | 32 |
| Pyruvate metabolism | 11 | 47 |
| Pyrimidine metabolism | 11 | 57 |
| Glycine and serine metabolism | 11 | 59 |
| Urea cycle | 10 | 28 |
| Histidine metabolism | 10 | 42 |
| Glycolysis | 9 | 23 |
| Beta-alanine metabolism | 9 | 34 |
| Aspartate metabolism | 9 | 35 |
| Glutathione metabolism | 8 | 20 |
| Transfer of acetyl groups into mitochondria | 8 | 22 |
| Fructose and mannose degradation | 8 | 31 |
| Amino sugar metabolism | 8 | 33 |
| Galactose metabolism | 8 | 38 |
| Glucose-alanine cycle | 7 | 13 |
| Mitochondrial electron transport chain | 7 | 19 |
| Phospholipid biosynthesis | 7 | 29 |
| Starch and sucrose metabolism | 7 | 31 |
| Nicotinate and nicotinamide metabolism | 7 | 35 |
| Methionine metabolism | 7 | 42 |
| Tryptophan metabolism | 7 | 59 |
| Nucleotide sugars metabolism | 6 | 20 |
| Carnitine synthesis | 6 | 22 |
| Plasmalogen synthesis | 6 | 26 |
| Pterine biosynthesis | 6 | 28 |
| Ammonia recycling | 6 | 31 |
| Sphingolipid metabolism | 6 | 40 |
| Propanoate metabolism | 6 | 42 |
| Valine leucine and isoleucine degradation | 6 | 59 |
| Glycerol phosphate shuttle | 5 | 11 |
| Cardiolipin biosynthesis | 5 | 11 |
| Phosphatidylcholine biosynthesis | 5 | 14 |
| Betaine metabolism | 5 | 21 |
| Glycerolipid metabolism | 5 | 25 |
| Cysteine metabolism | 5 | 26 |

| Pathway | Matched Entitles | Pathway Entitles of Experiment Type |
| --- | --- | --- |
| Pentose phosphate pathway | 5 | 29 |
| Folate metabolism | 5 | 29 |
| Lysine degradation | 5 | 30 |
| Tyrosine metabolism | 5 | 70 |
| De novo triacylglycerol biosynthesis | 4 | 9 |
| Malate aspartate shuttle | 4 | 10 |
| Phosphatidylethanolamine Biosynthesis | 4 | 12 |
| Alanine metabolism | 4 | 17 |
| Androstenedione metabolism | 4 | 24 |
| Estrone metabolism | 4 | 24 |
| Phytanic acid peroxisomal oxidation | 4 | 26 |
| Inositol metabolism | 4 | 30 |
| Androgen and estrogen metabolism | 4 | 33 |
| Retinol metabolism | 4 | 35 |
| Bile acid biosynthesis | 4 | 65 |
| Arachidonic acid metabolism | 4 | 67 |
| Ketone body metabolism | 3 | 13 |
| Butyrate metabolism | 3 | 19 |
| Lactose synthesis | 3 | 19 |
| Ethanol degradation | 3 | 19 |
| Threonine and 2-oxobutanoate degradation | 3 | 20 |
| Inositol phosphate metabolism | 3 | 24 |
| Selenoamino acid metabolism | 3 | 27 |
| Mitochondrial beta-oxidation of short chain saturated fatty acids | 3 | 27 |
| Mitochondrial beta-oxidation of longchain saturated fatty acids | 3 | 28 |
| Fatty acid elongation in mitochondria | 3 | 35 |
| Porphyrin metabolism | 3 | 40 |
| Fatty acid metabolism | 3 | 43 |
| Steroidogenesis | 3 | 43 |

**Table S3, related to Figure 1:** Complete list of pathways differentially regulated transcriptionally and metabolically in mouse SCLC cells in the *RPR2* model. For transcriptional regulation, adjusted P-value>0.05, n=4; for metabolic regulation, >3 metabolites.

| Pathway | P-value | Matched Entities |  |
| --- | --- | --- | --- |
| Oxidative stress and redox pathway | 1.25E-08 | 30 | 92 |
| Nicotine degradation II | 9.58E-05 | 11 | 49 |
| Glutathione-mediated detoxification | 5.95E-04 | 9 | 24 |
| Glutathione metabolism | 0.003645439 | 7 | 19 |
| Sphingolipid metabolism | 0.00390104 | 8 | 25 |
| Fatty acid Beta-oxidation II (core pathway) | 0.004422376 | 6 | 15 |
| Sphingolipid metabolism | 0.005170723 | 8 | 26 |
| Glutathione and one carbon metabolism | 0.006309387 | 9 | 44 |
| Cysteine biosynthesis III | 0.008129431 | 4 | 8 |
| Gamma-glutamyl cycle | 0.008129431 | 4 | 8 |
| Tryptophan metabolism | 0.008652042 | 10 | 44 |
| Arginine biosynthesis IV | 0.012455322 | 3 | 5 |
| Fatty acid beta-oxidation I | 0.015037244 | 7 | 24 |
| Ethanol degradation II | 0.020219266 | 4 | 11 |
| Bupropion degradation | 0.021756457 | 5 | 35 |
| Nicotine degradation III | 0.026358072 | 6 | 43 |
| D-myo-inositol (1,4,5)-trisphosphate biosynthesis | 0.026358072 | 6 | 22 |
| Superpathway of methionine degradation | 0.02873687 | 5 | 17 |
| Purine metabolism | 0.042260904 | 27 | 176 |
| Cyclic AMP biosynthesis | 0.0497073 | 11 | 58 |

**Table S4, related to Figure 3:** Target metabolites list for isotopologues extraction.

| Compound | Formula | Mass | CAS | RT (min) |
| --- | --- | --- | --- | --- |
| L-Serine | C<sub>3</sub>H<sub>7</sub>NO<sub>3</sub> | 105.04259 | 56-45-1 | 10.06 |
| Hypotaurine | C<sub>2</sub>H<sub>7</sub>NO<sub>2</sub>S | 109.01975 | 300-84-5 | 9.78 |
| L-Cysteine | C<sub>3</sub>H<sub>7</sub>NO<sub>2</sub>S | 121.01975 | 52-90-4 | 8.7 |
| Taurine | C<sub>2</sub>H<sub>7</sub>NO<sub>3</sub>S | 125.01466 | 107-35-7 | 8.41 |
| L-Glutamate | C<sub>5</sub>H<sub>9</sub>NO<sub>4</sub> | 147.05316 | 56-86-0 | 10.69 |
| Cystathionine | C<sub>7</sub>H<sub>14</sub>N<sub>2</sub>O<sub>4</sub>S | 222.06743 | 535-34-2 | 12.58 |
| L-Cystine | C<sub>6</sub>H<sub>12</sub>N<sub>2</sub>O<sub>4</sub>S<sub>2</sub> | 240.02385 | 56-89-3 | 12.62 |
| γ-Glu-Cys | C<sub>8</sub>H<sub>14</sub>N<sub>2</sub>O<sub>5</sub>S | 250.06234 | 636-58-8 | 11.12 |
| Glutathione | C<sub>10</sub>H<sub>17</sub>N<sub>3</sub>O<sub>6</sub>S | 307.08381 | 70-18-8 | 11.2 |
| Glutathione, oxidized | C<sub>20</sub>H<sub>32</sub>N<sub>6</sub>O<sub>12</sub>S<sub>2</sub> | 612.15196 | 27025-41-8 | 13.85 |

